# Mrs4 loss of function in fungi during adaptation to the cystic fibrosis lung

**DOI:** 10.1101/2023.04.05.535776

**Authors:** Daniel Murante, Elora G. Demers, Tania Kurbessoian, Marina Ruzic, Alix Ashare, Jason E. Stajich, Deborah A. Hogan

## Abstract

The genetic disease cystic fibrosis (CF) frequently leads to chronic lung infections by bacteria and fungi. We identified three individuals with CF with persistent lung infections dominated by *Clavispora* (*Candida*) *lusitaniae*. Whole genome sequencing analysis of multiple isolates from each infection found evidence for selection for mutants in the gene *MRS4* in all three distinct lung-associated populations. In each population, we found one or two unfixed, non-synonymous mutations in *MRS4* relative to the reference allele found in multiple environmental and clinical isolates including the type strain. Genetic and phenotypic analyses found that all evolved alleles led to loss of function of Mrs4, a mitochondrial iron transporter. RNA Seq analyses found that Mrs4 variants with decreased activity led to increased expression of genes involved in iron acquisition mechanisms in both low iron and replete iron conditions. Furthermore, surface iron reductase activity and intracellular iron was much higher in strains with Mrs4 loss of function variants. Parallel studies found that a subpopulation of a CF-associated *Exophiala dermatiditis* infection also had a non-synonymous loss of function mutation in *MRS4.* Together, these data suggest that *MRS4* mutations may be beneficial during chronic CF lung infections in diverse fungi perhaps for the purposes of adaptation to an iron restricted environment with chronic infections.

## Introduction

Evolution of pathogens in infections can lead to the rise of isolates with increased resistance to host defenses or drugs, improved fitness, or enhanced access to nutrients. An understanding of pathoadaptive mutations may improve therapies and treatments. The repeated rise of specific mutations in bacteria and fungi associated with chronic infections has been particularly well-documented in the context of lung infections in people with cystic fibrosis (CF). The genetic mutations that cause CF lead to chronic infections. Several studies have shown that nutritional immunity, mediated by innate immune effectors such as calprotectin which restrict access to metals, force microbes to employ diverse strategies to acquire essential nutrients such as iron and zinc (1–3).

Many of the mutations that repeatedly arise in CF-related bacterial and fungal pathogens are in regulators (e.g. *lasR* and *mucA* in *Pseudomonas aeruginosa*, *agr* in *Staphylococcus aureus*). Analysis of *Candida* CF isolates has found similar regulatory mutations. A study of *Candida albicans* CF infection identified six instances of loss-of-function (LOF) mutations in *NRG1,* which encodes a repressor of filamentation; *nrg1* LOF mutants are resistant to the suppression of filamentation by the frequently coinfecting bacterium *Pseudomonas aeruginosa* (4). We previously published that a single *Clavispora* (*Candida*) *lusitaniae* infection, with no detectable co-infecting bacteria, had numerous activating and subsequent suppressing mutations in *MRR1* (5) that lead to heterogenous resistance to the antifungal fluconazole, the toxic metabolite methylglyoxal, and *P. aeruginosa* toxins. Longitudinal collections of *Aspergillus fumigatus* isolates from a single individual with CF showed acquired mutations that lead to HOG pathway hyperactivation and improved fitness in the presence of oxidative and osmotic stress (6).

In this work, we describe a locus in *C. lusitaniae* (7) that was independently mutated in three separate subjects with CF. *C. lusitaniae*, a haploid member of the CTG clade within the Saccharomycetaceae family, is known to readily develop Amphotericin B resistance (8) and can develop resistance to caspofungin and azoles (9, 10). *C. lusitaniae* has been reported in association with plant and food products, and is a less commonly found to be an abundant member of microbiome communities. Notably, *C. lusitaniae* is closely related to *Candida auris* which is also known for the repeated development of multi-drug resistance, and is a critical threat on the WHO priority pathogens list (11, 12).

*C. lusitaniae* evolution in the CF lung may provide an opportunity to study fungal adaptation to host environments. We found non-synonymous mutations in *MRS4* arose independently in three different CF lung infections. Mrs4 is a high affinity inner mitochondrial membrane iron transporter that brings iron into the inner lumen. In other fungi, Mrs4-mediated iron transport is necessary for robust growth in low iron environments, resistance to Cons and menadione, and its function supports the synthesis of iron-sulfur clusters for incorporation into diverse enzymes and regulators (13–18). Previous studies found that *MRS4* deletion significantly reduces virulence of *C. albicans* in a murine model for systemic candidiasis (19). We found that each of the Mrs4 variants had decreased function. RNA seq analysis of isolates with different *MRS4* alleles in iron replete and iron restricted conditions demonstrated that Mrs4 LOF led to significant increases in expression of multiple iron uptake methods. Furthermore, strains with *MRS4* LOF alleles demonstrated increased accumulation of intracellular iron. A LOF mutation in *MRS4* was also found in CF infection isolates of *Exophiala dermatitidis*. Taken together, these data highlight that in two distinct environmental fungi, chronic infection leads to the selection for Mrs4 variants with decreased function and an increased capacity for iron uptake. Future studies will determine if these host-adapted strains have new vulnerabilities that can be exploited therapeutically.

## Results

### *MRS4* mutations are found in *C. lusitaniae* from CF lung infections

Analysis of the microbiota in bronchoalveolar lavage (BAL) fluids collected at Dartmouth Health found three subjects CF with lung infections dominated by *C. lusitaniae* ((7) and in a manuscript in preparation). We sequenced the genomes of 12-20 isolates from each population (see Methods section for Accession numbers). To identify mutations that likely arose during infection, non-synonymous single nucleotide polymorphisms (SNPs) that were not fixed within the population from each individual were determined (**Table S1**). Nineteen genes had two alleles with non-synonymous differences in more than one population, defined as isolates from a single subject at a single time point, and only one gene had alleles with non-synonymous differences within in all three populations: *CLUG_02526* (**Figure 1A**). *CLUG_02526* encodes an amino acid sequence with 71% identity with *C. albicans* SC5314 Mrs4 (19), and 51% with *Saccharomyces cerevisiae* S288C Mrs3 and Mrs4 (**Figure S1**). Mrs4 functions as high affinity mitochondrial iron importers in both species (16, 20). Due to the high percent sequence identity and our phenotypic characterization of *CLUG_02526* deletion mutants (described below), we will heretofore refer to *CLUG_02526* as *MRS4*.

**Figure 1.**
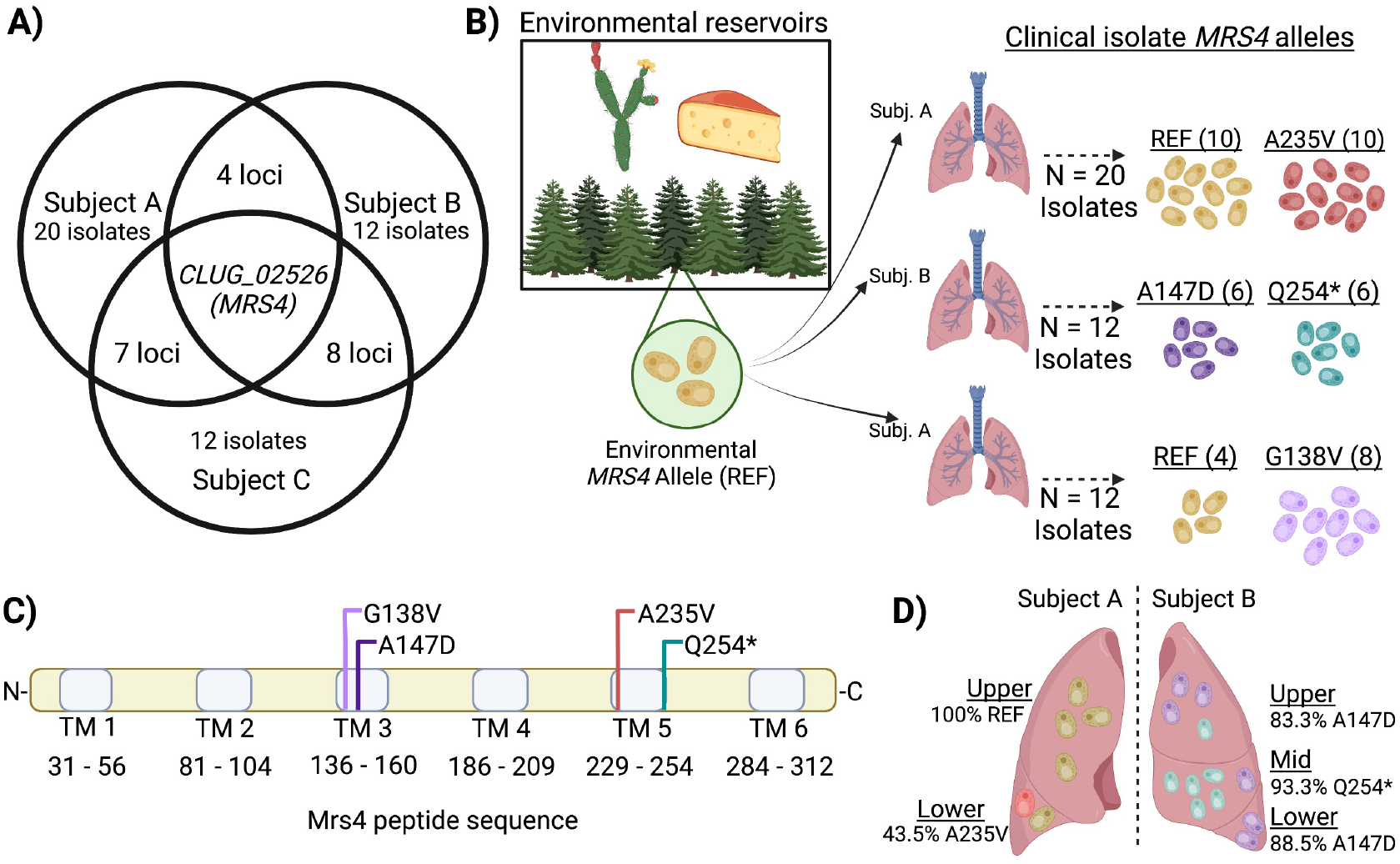
Non-synonymous SNPs in *MRS4* were found in whole genome sequence data from 12-20 *C. lusitaniae* isolates from each of three subjects with chronic CF infections. **A)** Analysis of loci that were heterogeneous in three *C. lusitaniae* populations in three separate individuals (Subjects A, B, and C) found that only *CLUG_02526* (*MRS4*) had subpopulations with non-synonymous substitutions in all three infections. **B)** Two *MRS4* alleles were detected in each population. “REF” indicates the *MRS4* sequence in environmental and acute infection isolates of *C. lusitaniae.* **C)** The *MRS4* sequence encodes a barrel-structure iron transporter on the inner mitochondrial membrane; the protein is 318 amino acids long and comprised of six transmembrane alpha-helices denoted by light blue bars. Each mutation is predicted to disrupt or truncate one of these transmembrane domains (see SUSPECT analysis, Fig S3). **D)** Pooled sequencing was performed on isolates from bronchoalveolar lavage fluid taken from specific lobes of subjects A and B. The relative abundances of *MRS4* alleles was quantified by analysis of individual reads.

To compare the *MRS4* sequences from CF *C. lusitaniae* isolates to *MRS4* sequences to a broader collection of *C. lusitaniae* strains, we also analyzed *MRS4* in ten *C. lusitaniae* isolates from diverse clinical and environmental sources. We found that the predicted Mrs4 amino acid sequences were identical and the genes differed by only a small number of synonymous SNPS that varied among alleles (**Figure S2**). The conserved Mrs4 amino acid sequence will be referred to as the “reference” or Mrs4^REF^ sequence. Isolates with *MRS4* alleles that encoded the Mrs4^REF^ sequence were found in both Subject A and Subject C (**Figure 1B**). In addition to the reference allele, Subject A and Subject C populations each had isolates with mutant alleles that differed by single non-synonymous SNPs, and encoded *MRS4^A235V^*and *MRS4^G138V^*, respectively (**Figure 1B**). Subject B isolates carried one of two mutant alleles that encoded *MRS4^A147D^* and *MRS4^Q254*^* (**Figure 1B**), suggesting that two independent *MRS4* mutant lineages arose within that population. The Mrs4 substitutions or terminations occurred in predicted transmembrane alpha helices of the protein (**Figure 1C**) and occurred at residues that were conserved across diverse species (**Figure S1**). Each *MRS4* mutation found in the CF *C. lusitaniae* isolates had a high likelihood of affecting function based on the SuSPECT analysis method which estimates the probability for single amino acid variants to impact phenotype (**Figure S3**) (21).

### Spatial and longitudinal analyses show that *MRS4* alleles may have arisen independently in different regions of the lung and that Mrs4 LOF isolates persisted over time and across compartment

Using whole genome sequence data for pools of 75-96 isolates from the upper and lower lobes of the right lung of Subject A and the upper, middle and lower lobes of the left lung of Subject B, we determined the fraction of reads that encoded the *MRS4* SNPs described above within each pool. For the pooled isolates of the upper lobe of Subject A, the reads contained only the *MRS4^REF^*allele, while ∼43% of sequenced population from the lower lobe had the *MRS4^A235V^*allele (**Figure 1D**). Both alleles were detectable in the upper, middle, and lower lobe isolate pools of Subject B, with the *MRS4^A147D^* allele present at a higher percentage in the upper and lower lobe (∼83 and 89%, respectively), while *MRS4^Q254*^*was found in ∼93% of reads in the middle lobe. Sequence analysis of four sputum isolates from a sample donated by subject A, collected ∼1 year after the BAL isolates were recovered, found two isolates with *MRS4^REF^* and two isolates with *MRS4^A235V^* suggesting that the MRS4 variant-containing isolates persisted over time. Similarly, we obtained 6 respiratory sputum and stool isolates from Subject B more than one year after the initial isolation and amplified and sequenced the *MRS4* allele. We found that they all contained the *MRS4^A147D^* allele. These longitudinal isolates indicate that the *MRS4* mutations persisted over time and possibly in multiple compartments, and thus were not transiently present at the time of isolation.

### *MRS4* variants confer LOF phenotypes

In other fungi, the deletion of *MRS4* impairs growth in low iron media or the presence of an iron chelator (17, 20, 22). To determine the activity of Mrs4 variants in the *C. lusitaniae* CF isolates, we first constructed an *mrs4*Δ derivative of Subject B isolate B_L01, which had an *MRS4^Q254*^* allele, then complemented back either the native B_L01 *MRS4^Q254*^* allele or *MRS4^REF^* at the native locus (**Figure 2A**). We chose the *MRS4* allele from strain ATCC 42720 as the source of the *MRS4^REF^* sequence. We observed that all four isogenic strains (B_L01 parental isolate, the *mrs4*Δ mutant, and the *mrs4*Δ mutant complemented with *MRS4^REF^*or *MRS4^Q254*^*) reached similar yields in YPD (**Figure 2B**). Yields were reduced in YPD amended with a high-affinity ferrous iron chelator bathophenanthroline disulfonate (BPS), and we observed differences dependent on *MRS4* allele (**Figure 2B**). Deletion of *mrs4* in the B_L01 background reduced final yield. Complementation with the *MRS4^REF^* allele restored growth to a significantly greater degree than the parent strain, or to the B_L01 *mrs4Δ* complemented with the native allele. These results indicate that the truncated variant has decreased function relative to the reference allele, but that the truncated variant may retain some function.

**Figure 2.**
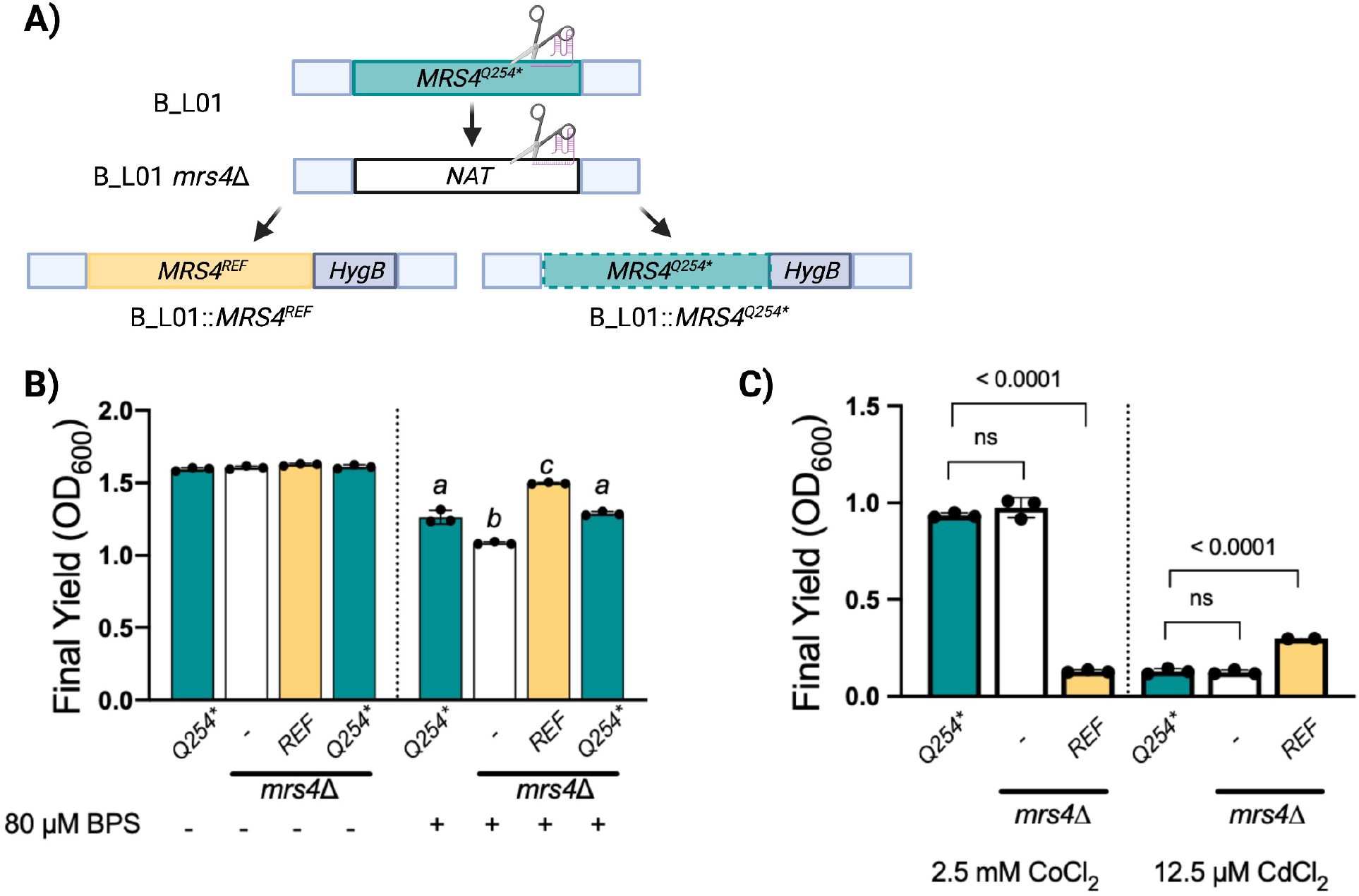
Mrs4^Q254*^ confers loss of function in *C. lusitaniae*. **A)** An *mrs4Δ* mutant and *mrs4*Δ mutants complemented with *MRS4^Q254*^* or the *MRS4^REF^* were constructed in the B_L01 clinical isolate background **B)** Strains were assessed for growth in a 96-well plate after 24 h at 37° in YPD or YPD with 80 µM BPS iron chelator. Columns labelled with *a* are non-significantly different from each other, and are significantly different from columns labelled with *b* and *c.* **C)** Indicated strains were grown for 24 h at 37° in YPD supplemented with 2.5 mM CoCl_2_ (left) and 12.5µM CdCl_2_ (right). There were at least three replicates per sample. Indicated p-values are from a one-way ANOVA with Tukey’s post-hoc correction, ns, not significant.

Deletion mutants lacking *MRS4* in *S. cerevisiae* and *C. albicans* have altered metal sensitivities such that mutants are more resistant to cobalt and more sensitive to cadmium when compared to their Mrs4+ counterparts (17, 20) due to activation of transcription factors such as Aft1 in *S. cerevisiae* (15). We found that cadmium and cobalt affected the growth of *C. lusitaniae* in an *MRS4* allele dependent manner. The B_L01 *mrs4*Δ derivative and the *mrs4*Δ complemented with the allele encoding Mrs4^Q254*\**^ were more resistant to cobalt than the *mrs4Δ* mutant complemented with *MRS4^REF^* (**Figure 2C**). Conversely, the *mrs4*Δ strain complemented with the Mrs4^REF^ variant was more resistant to cadmium than the *mrs4*Δ mutant or the mutant complemented with Mrs4^Q254*\**^. Together, these data further suggest that the Mrs4^Q254*\**^ variant is less functional than the reference allele.

To characterize the levels of function for the other three Mrs4 variants found in the CF clinical *C. lusitaniae* isolates, we created *mrs4*Δ derivatives of isolates A_U05 (*MRS4^A235V^),* B_L04 (*MRS4^A147D^*), and C_M06 (*MRS4^G138V^*) which were then complemented with the *MRS4^REF^*allele. In each case, complementation with the functional *MRS4^REF^*allele made cells more sensitive to cobalt (**Figure 3A**) as was the case for strain B_L01. Similarly, replacement of *MRS4* with the *MRS4*^REF^ allele increased cadmium sensitivity in B_L04 (*MRS4^A147D^*), and C_M06 (*MRS4^G138V^*) (**Figure 3B**). Unexpectedly, the A_U05 (*MRS4^A235V^*) isolate had greater resistance to cadmium than the same background with the *MRS4^REF^* allele, perhaps due to other genetic differences. When we expressed *MRS4^A235V^* in B_L01 *mrs4*Δ, the resultant strain was more sensitive to cadmium than the isogenic strain with *MRS4^REF^* supporting the conclusion that the Mrs^A235V^ variant had low or no activity (**Figure S4**). We also made an *mrs4*Δ mutation in an outgroup *C. lusitaniae* strain RSY284 (DH2383) (23) with a native *MRS4*^REF^ allele, and confirmed that the mutant had the expected resistance to cobalt and sensitivity to cadmium (**Figure S5A**). Together, these data suggest that LOF mutations in *MRS4* arose four independent times across three chronic CF infections indicating possible selection for phenotypes associated with loss of Mrs4 function.

**Figure 3.**
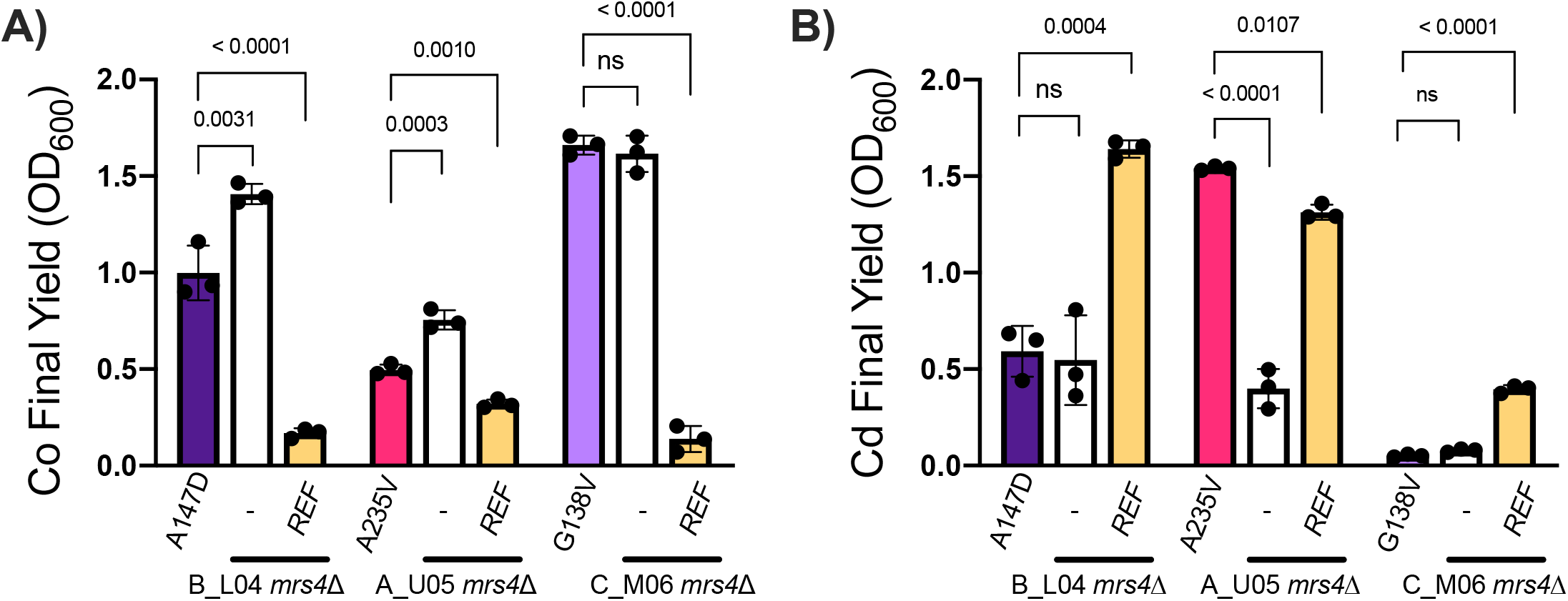
*MRS4* mutations in each clinical population demonstrate LOF phenotypes. Representative parent isolates of each mutation from Subject B (B_L04, *MRS4^A147D^*), Subject A (A_U05, *MRS4^A235V^*), and Subject C, (C_M06, *MRS4^G138V^*) and their *mrs4Δ* derivatives that were then complemented with the *MRS4^REF^* allele, were grown in YPD supplemented with **A)** 2.5 mM CoCl_3_ and **B)** 12.5 µM CdCl_2_. Data represent the endpoint OD600 measured by a Synergy Neo2 plate reader after 24 h of growth at 37°. Indicated p-values are from one-way ANOVA with Tukey’s post-hoc, ns, not significant.

#### Characterization of *MRS4* growth phenotypes

Mitochondrial metabolism is highly dependent on the availability of metals such as iron. To assess whether the *MRS4* loss-of-function alleles affected mitochondrial activity, we examined the growth of isogenic strains with either *MRS4*^REF^ or Mrs4^Q254*^ in medium with either glucose (which can be fermented) or glycerol as the major carbon source. In both yeast extract peptone medium or yeast nitrogen base media with 2% glucose, there was a slightly faster growth for the strain with Mrs4^REF^ compared to the *mrs4* null mutant or strains with Mrs4^Q254*^ but the differences were not significant (**Figure 4A and C**). Similar results were obtained when grown in medium with glycerol as the dominant or sole carbon source (**Figure 4B and D**). These results suggested that there was no major defect in respiratory metabolism.

**Figure 4.**
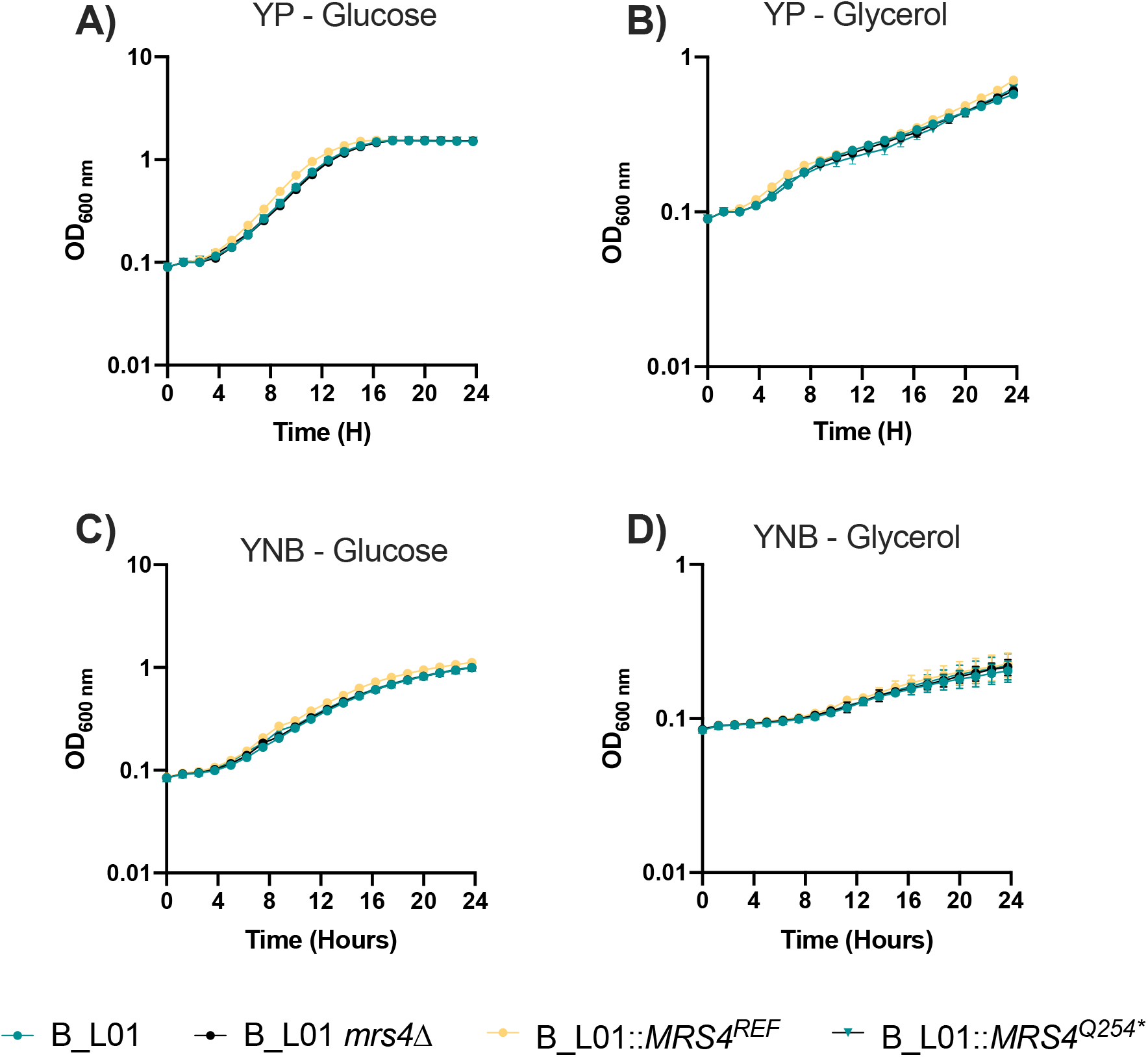
Lack of MRS4 growth phenotypes in complex and minimal media with glycerol and glucose. Cells from exponential phase cultures of B_L01, its *mrs4*Δ derivative, and *mrs4*Δ mutants complemented with *MRS4^Q254*^* or the *MRS4^REF^* were inoculated into a 96-well plate and grown for 24 h at 37° in YP medium supplemented with **A)** 2% glucose or **B)** 2% glycerol or in YNB defined medium without amino acids supplemented with **C)** 2% glucose or **(D)** 2% glycerol. OD_600_ was measured over time using a Synergy Neo2 plate reader.

As a more sensitive indicator of changes in levels of respiration and fermentation, we quantified glucose consumption relative to fermentation product production by HPLC. Analysis of supernatants from cultures of the *mrs4*Δ + *MRS4^REF^* strain grown in medium with glucose as the sole carbon source found that the dominant fermentation product was acetate followed by ethanol; glycerol was not detected. Under the same conditions, *C. albicans* strain SC5314 produced mainly ethanol with low levels of acetate and glycerol. Normalized to moles of carbon, *C. lusitaniae* converted more than twice as much glucose to fermentation products than *C. albicans* (**Table S2**). When the mass balance of glucose consumed and fermentation products made for *C. lusitaniae mrs4*Δ + *MRS4^Q254*^* and *mrs4*Δ + *MRS4^REF^* were compared, we found no significant differences suggesting comparable levels of respiration and fermentation (**Table S2**). We also constructed a *C. albicans mrs4Δ/Δ* homozygous mutant and found no significant difference in fraction of carbon used for fermentation when the SC5314 wild type was compared to the *mrs4Δ/Δ* homozygous mutant (**Table S2**).

To further assess metabolic differences associated with Mrs4 LOF, we analyzed growth of B_L01 parent and *mrs4Δ*::*MRS4^REF^*on different sole carbon sources using the Biolog™ carbon utilization phenotype microarray plates. We inoculated equal concentrations of each strain in YNB minimal medium and observed growth over the course of 48 h. Across the 192 carbon sources tested, there were no differences in growth that were confirmed in secondary analysis (**Supplementary dataset 1**). Analysis of B_L01 parent and B_L01::*MRS4^REF^* did not show any differences in minimal inhibitory concentrations for commonly used antifungals including fluconazole and amphotericin B (data not shown). Consistent with published studies in *Cryptococcus neoformans* (24), in *C. lusitaniae* DH2383 and B_L01, functional Mrs4 was necessary for full H_2_O_2_ resistance (**Figure S5**).

### *MRS4* LOF impacts expression of iron homeostasis regulators and metal storage

To gain insight into how Mrs4 LOF affected *C. lusitaniae*, we performed a transcriptomics analysis of B_L01 *mrs*Δ with *MRS4^REF^* and *MRS4^Q254*^*. Because cells with defective Mrs4 (e.g. *MRS4^Q254*^*) showed reduced growth on medium with chelator (**Figure 2B**), we performed an RNA-seq analysis of cells from mid-exponential phase cultures growing in YPD and parallel cultures that received a 1 h exposure to the iron chelator BPS (**Figure 5A** for experimental scheme). The short exposure to BPS was used to limit the pleiotropic effects associated with growth differences. Comparison of the strains with *MRS4^REF^*without or with exposure to iron chelator found eleven genes that had a fold difference greater than log_2_ 1 and a false discovery corrected p-value less than 0.05 (**Figure 5B** and **Supplementary dataset S2**). Among the genes that were differentially expressed were four cell surface high affinity ferric iron uptake genes (*FTR1*, *FRE9*, *FRE10*, and *FET31*) and genes that encode regulators involved in iron utilization *SFU1* and *HAP43* (**Figure 5B**) which are known to be transcriptionally regulated in response to iron limitation response in other *Candida* species (25, 26).

**Figure 5.**
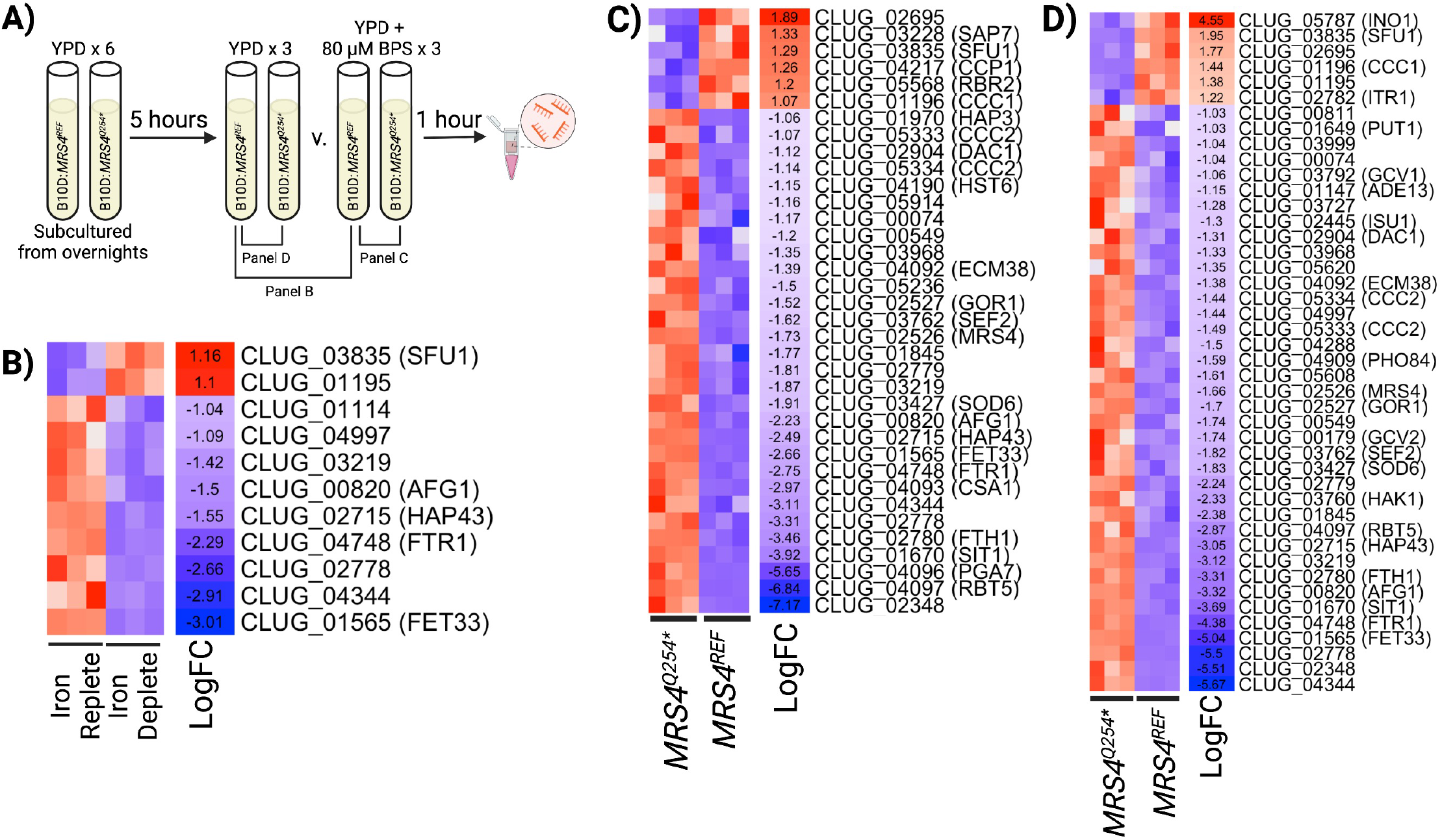
Loss of Mrs4 function leads to increased expression of iron acquisition genes. **A)** Design of RNA-seq sample preparation. Sextuplicate cultures of B_L01 *mrs4*Δ complemented with either *REF* or *Q254* MRS4* alleles were grown overnight, then sub-cultured into YPD and grown for 5 h. Cultures were grown for an additional hour with either 80µM BPS or vehicle prior to RNA isolation. Gene expression heatmaps of differentially expressed genes (P < 0.05 and a log_2_ fold-change ≥|1|) in a comparison between **B)** the B_L01::*MRS4^REF^*strain grown in YPD (iron replete) or YPD with BPS (iron deplete) **C)** B_L01::*MRS4^REF^*and B_L01::*MRS4^Q254*^* grown in YPD with BPS, and **D)** B_L01::*MRS4^REF^* and B_L01::*MRS4^Q254*^* grown in YPD.

Comparison of the transcriptomes of B_L01 *mrs*Δ+*MRS4^REF^* to B_L01 *mrs4*Δ+*MRS4^Q254*^*after chelator exposure found that the same ferric reductases that were induced upon exposure to chelator in the Mrs4^REF^ strain were higher in cells with Mrs4^Q254*^ (**Figure 5C**). We again found differential expression of the *SFU1* and *HAP43* regulators, and also found greater fold induction of the gene encoding pro-iron acquisition transcription factor *SEF1* in the *mrs4*Δ+*MRS4^Q254*^* relative to *mrs*Δ+*MRS4^REF^* with chelator. In addition, we found differential expression of putative orthologs of iron uptake genes (*SIT1*, *CSA1*) and other metal transporters (*CCC1*, *CCC2*) (27) when Mrs4 function was low. *MRS4* transcript levels were log_2_ 1.7 fold higher in the *mrs4*Δ+*MRS4Q254*\* strain, and the *MRS4*-adjacent gene *GOR1*, predicted to encode a glyoxylate reductase, was also significantly higher in the Mrs4^Q254*^ bearing strain.

We also compared transcriptomes of *mrs*Δ+*MRS4^REF^* and *mrs4*Δ+*MRS4^Q254*^* in control conditions without chelator. Thirty-seven genes were differentially expressed across the two strains in the presence of chelator (**Figure 5C**), and 27 of them were still differentially expressed in its absence (**Figure 5D**). Ferric reductase encoding genes and their regulators (e.g. *HAP43*) were again differentially expressed, and the fold difference between strains was higher than in the chelator condition (**Figure 5C** and **D**). An independent experiment, using qRT-PCR analysis of RNA from *mrs*Δ+*MRS4^REF^* and *mrs4*Δ+*MRS4^Q254*^*also found transcript levels of *HAP43* and *FRT1* to be significantly higher in both iron replete and iron chelated conditions (**Figure S6**). We also observed that *mrs*Δ+*MRS4^REF^* had ∼4.5 fold higher levels of *INO1*, the ortholog of inositol-3-phosphate synthase, than *mrs4*Δ+*MRS4^Q254*^*in YPD (**Figure 5D**).

### Loss of Mrs4 function leads to higher surface ferric reductase activity

We sought to determine if the higher levels of in transcripts encoding cell surface ferric reductases in *MRS4^Q254*^* strains, even in iron-replete conditions, translated into higher levels of iron acquisition activity. To do so, we utilized tetrazolium chloride (TTC), a substrate for ferric reductases (28). To avoid complications associated with growth inhibition by TTC, we grew colonies on YNB-glycerol agar for 24 h, then overlaid with molten 1% agar containing TTC (**Figure 6A**). Upon TTC reduction by ferric reductases, an insoluble red pigment forms. After ten minutes, there was a strong red coloration associated with colonies formed by *MRS4^Q254^*\* and the *mrs4*Δ strains, but not the *MRS4^REF^*strain (**Figure 6A**). The TTC reduction phenotype was abrogated by the addition of excess ferric iron in the agar overlay (**Figure 6A**). These data suggest greater ferric reductase activity on the cell surface when Mrs4 activity is low. Similar results though to a lesser degree were observed on YPD medium (**Figure 6B**).

**Figure 6.**
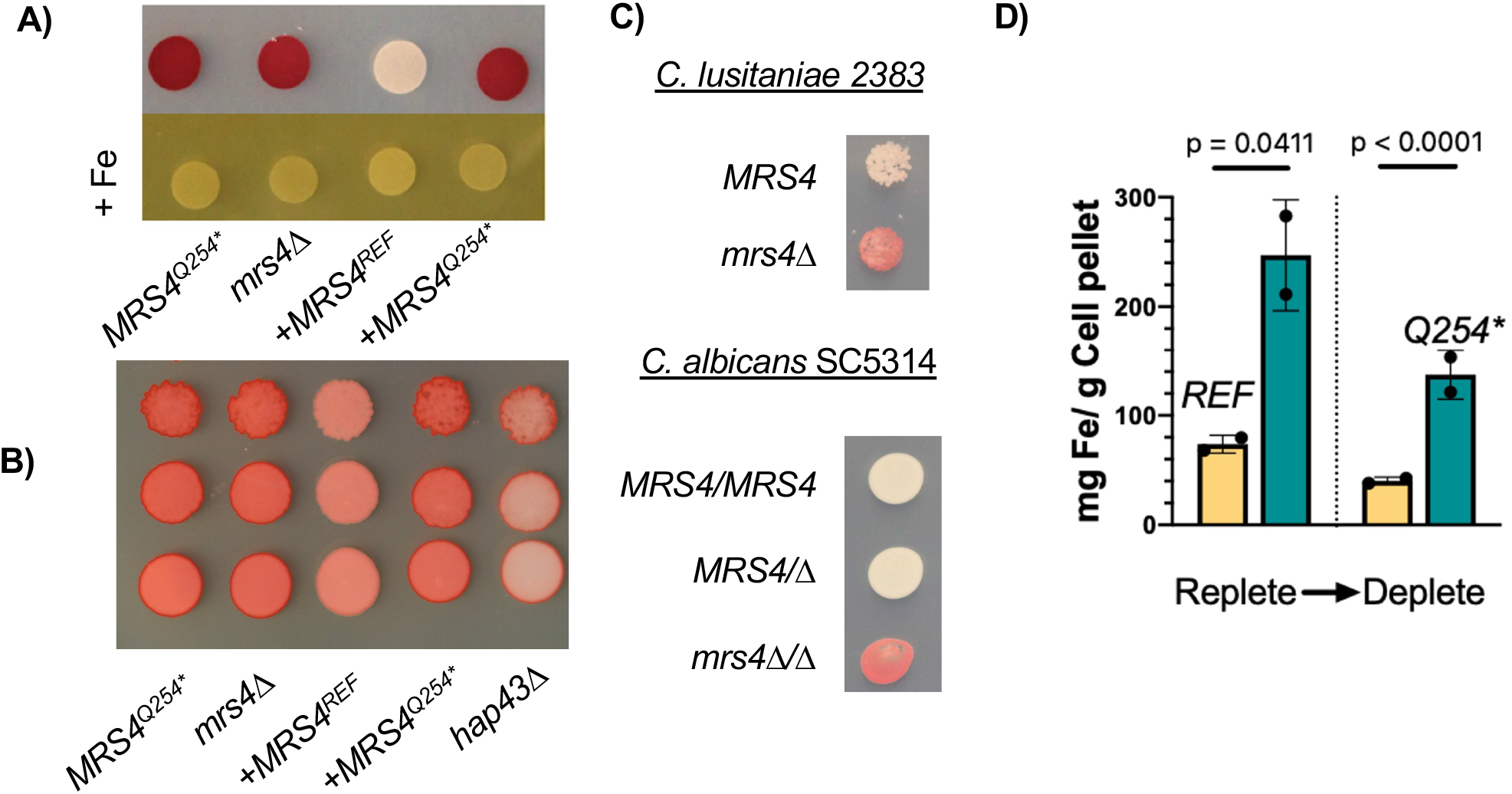
Decreased Mrs4 function increases ferric reductase activity and intracellular iron content. **A)** B_L01 derived strains *mrs4Δ, mrs4Δ::MRS4^REF^, mrs4Δ::MRS4^Q254*^*were spotted on YNB-glycerol plates. Plates were incubated for 24 h at 37°C. Each plate was overlayed with a 10 ml solution of 1 mg/ml tetrazolium chloride (TTC) and incubated for 5 min prior to imaging. Red pigmentation indicated represents greater levels of ferric iron reduction. Inclusion of 10 mM of FeCl_3_ (+Fe) as a competitor eliminates TTC reduction. 24 **B)** B_L01 derived strains *mrs4Δ, mrs4Δ::MRS4^REF^, mrs4Δ::MRS4^Q254*^* and *hap43*Δ were serially diluted from 1 OD and spotted on YPD plates, then allowed to grow for 24 hours at 37**°** C then analyzed as in panel A. **C)** *C. lusitaniae* DH2383 and its *mrs4Δ* mutant and *C. albicans* SC5314 with single and double knockouts of *mrs4* were analyzed for surface ferric reductase activity on YNB-glycerol. **D)** B_L01 *mrs4Δ* strains complemented with *MRS4^REF^* and *MRS4^Q254*^* alleles were grown in YPD or YPD + 80 µM BPS as outlined in Fig. 5A. Whole cell iron was quantified using ICP-MS. Data represents the averages of three technical replicates for two experiments done on separate days. Indicated p-values are from Student’s t-tests.

Previously studies showed that decreased mitochondrial iron levels induced the activity of *C. lusitaniae* transcription factors by modulating cytosolic Fe-S-containing regulators. We predicted that Hap43 (within the Hap43-Sfu1-Sef1 transcription factor network) was part of this response that led to increased expression of iron uptake genes. Thus, we analyzed a *hap43*Δ mutant in the B_L01 background with the defective Mrs4^Q254*^ variant, and found that Hap43 indeed contributed to the increased levels of iron reductase activity created by Mrs4 LOF (**Figure 6B**). We also found that surface ferric reductase activity was higher in the *mrs4*Δ strains if *C. lusitaniae* strain 2383 and *C. albicans* SC5314 *mrs4*Δ/Δ. In *C. albicans*, both copies of *MRS4* to be deleted for manifestation of the increased surface ferric reductase phenotype.

### Mrs4 LOF leads to the accumulation of intracellular iron

To evaluate the consequences of differential expression of iron acquisition genes in strains with and without Mrs4 function, we analyzed the concentrations of cellular iron by ICP-MS. In iron replete YPD medium, the strain *mrs4*Δ+ *MRS4^Q254*^* had significantly higher levels of intracellular iron than *mrs4*Δ::*MRS4^REF^*(**Figure 6D**). In cells subjected to chelator treatment for 1 h prior to harvest, intracellular iron was lower than in the untreated cells, but still significantly higher in the *mrs4*Δ::*MRS4^Q254*^* strain. These data suggest that the observed increase in iron acquisition gene transcripts was concomitant with higher intracellular iron which may be advantageous *in vivo* and may explain the repeated LOF of Mrs4 in clinical *C. lusitaniae* populations.

### *MRS4* mutations in other fungi of clinical interest

In light of variation in *MRS4* in three different *C. lusitaniae* populations, we investigated the consequences of two other naturally-occurring *MRS4* allelic variations that we observed in other species. First, we assessed the activity of Mrs4 variants in *Candida auris,* a fungus that emerged within the past forty years (29) and is closely related to *C. lusitaniae*. Multidrug resistant strains that caused localized outbreaks emerged independently within genetically distinct clades. Analysis of *Candida auris MRS4* alleles within and between clades, available from published sequences (30), found that the encoded Mrs4 sequences were identical, with one exception. Clade I strains differed from strains in the other clades in that the *MRS4* allele encoded Mrs4^31V^ while others encoded Mrs4^31A^. This difference was confirmed by Sanger sequencing. To assess the activity of these two Mrs4 variants, we expressed them in a *C. lusitaniae mrs4*Δ mutant. Both *C. auris* Mrs4^31A^ and Mrs4^31V^ alleles were able to complement *mrs4*Δ for growth with iron chelator and cadmium sensitivity to a similar extent as the functional *C. lusitaniae* Mrs4^REF^ protein (**Figure S7**), indicating that both alleles were functional.

The second analysis of Mrs4 variants was in the black yeast *Exophiala dermatiditis*. We had previously identified a CF infection predominated by *E. dermatiditis.* We performed whole genome sequencing of twenty-three isolates and found a subpopulation of isolates with a non-synonymous SNP in the *MRS4* (**Figure 7A**) (31). Subsequent Sanger sequencing confirmed that seven of the twenty three isolates carried an *MRS4* ortholog with a sequence identical to the *E. dermatiditis* type strain, NIH8656 (referred to here as encoding Mrs4^40E-REF^). The other sixteen sequenced isolates had a variant *MRS4* with an E40G substitution. To characterize the *MRS4* alleles in these *E. dermatitidis* isolates, both were synthesized and heterologously expressed in *C. lusitaniae mrs4*Δ and assessed for function relative to the *C. lusitaniae MRS4* alleles. The *E. dermatiditis* alleles (*EdMRS4^40E-REF^* and *EdMRS4^40G^*) were codon optimized for *Candida* spp. and the spliced introns were removed. The *EdMRS4* alleles were introduced at the native *MRS4* site and expressed under the control of the *C. lusitaniae MRS4* promoter.

**Figure 7.**
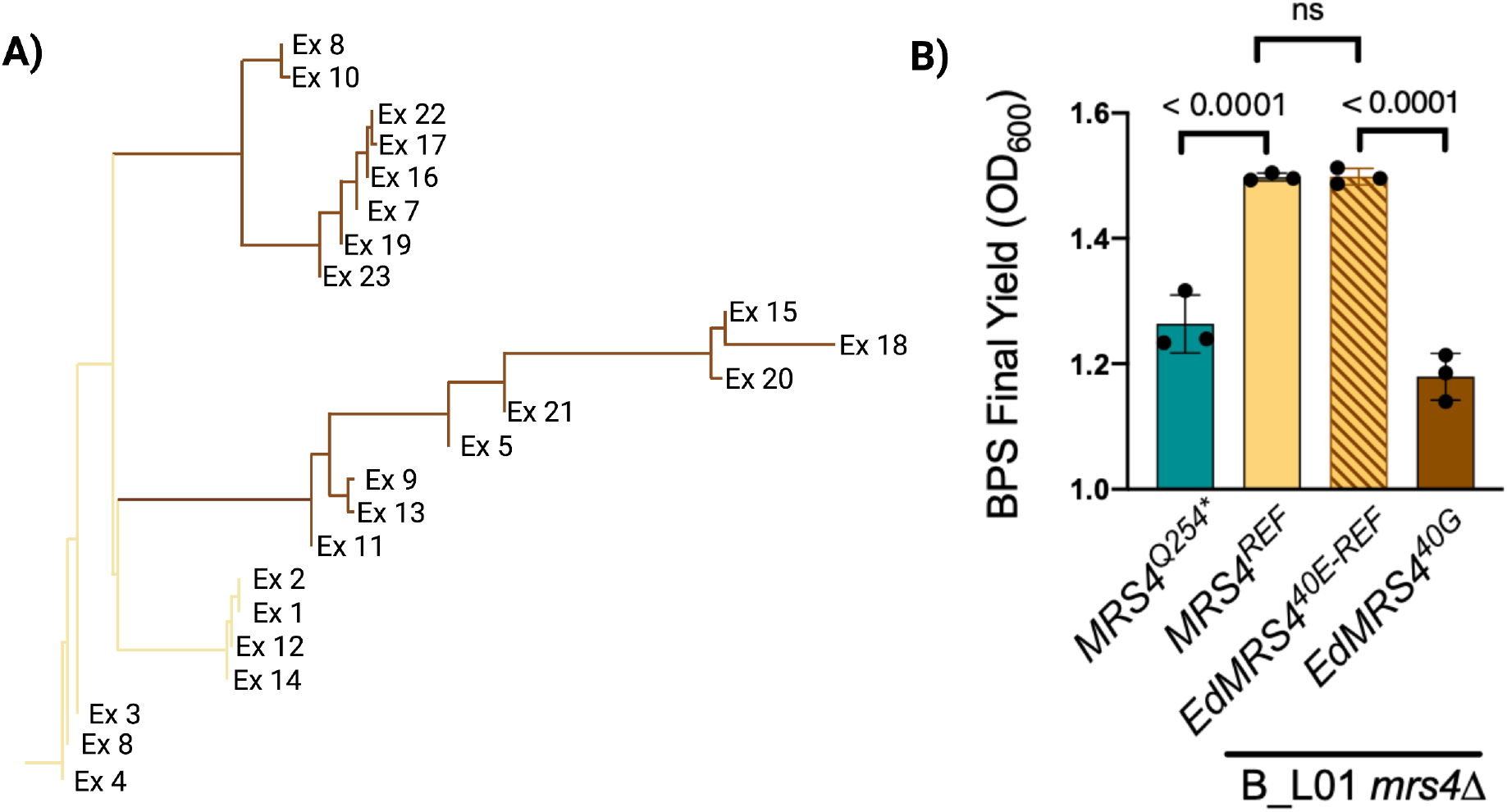
An Mrs4 loss-of-function subpopulation also emerged in *Exophiala dermatiditis* during a chronic CF lung infection. **A)** Two alleles of *MRS4* were found in *E. dermatitidis* isolates from in a single chronic CF lung infection. Of the 23 isolates sequenced, seven genomes encoded the reference Mrs4^40E^ (*E.d. REF*), which is identical to previously sequenced *E. dermitiditis* strains, and sixteen isolates encoded an Mrs4^40G^ variant (*E40G*). **B)** *C. lusitaniae* B_L01 *mrs4*Δ strains complemented with the two *E. dermatitidis MRS4* alleles grown in YPD with 80 µM BPS for 24 h. *C. lusitaniae* B_L01 *mrs4*Δ expressing functional *MRS4^REF^* or *MRS4^Q254*^* were included for comparison. Indicated p-value are from a one-way ANOVA with Tukey’s post-hoc, ns, not significant.

Complementation of the *mrs4*Δ strain with Mrs4^40E-REF^ fully complemented the *mrs4*Δ mutant to the levels of *C. lusitaniae MRS4*^REF^. In the presence of the BPS iron chelator, the *mrs4*Δ strain with the *E. dermatiditis* Mrs4^40G^ variant had significantly reduced growth compared to a strain with *E. dermatiditis* Mrs4^40E^, which fully restored the Mrs4 function to levels observed for the *C. lusitaniae* reference gene (**Figure 7B**). This indicates that a Mrs4 LOF mutation also arose in a population of *E. dermatitidis* during a CF lung infection. The repeated occurrence of *MRS4* mutations in fungal CF infections strongly suggests a selective benefit for *MRS4* LOF mutations in the CF lung.

## Discussion

In this work, we showed the repeated loss of Mrs4 activity across two fungal species, *C. lusitaniae* and *E. dermatiditis*, in the context of chronic CF lung infections. Each acquired non-synonymous mutations that resulted in reduced or loss of Mrs4 function, which led to defects in iron import into the mitochondrial inner lumen and increased expression and activity of iron acquiring pathways. The selection for *MRS4* LOF mutations in chronic lung infection may inform future studies on the mechanism of fungal persistence within the host and resistance to therapeutic strategies. The emergence of Mrs4 LOF in two diverged species of Ascomycota, both environmental fungi which colonized chronic CF lung infections, highlights the possibility that mutations in *MRS4* may be important for the shift to commensal colonizing yeast. Work by Kim et al. (4) found *C. albicans NRG1* LOF mutations in isolates from different individuals with CF suggesting that inactivating mutations in this locus increased fitness. Interestingly, mutants in Nrg1 have increased expression of iron uptake genes (32).

In other species, mutation of Mrs4 leads impairs the mitochondrial synthesis of Fe-S clusters, and Fe-S cluster levels modulate the iron starvation response through their insertion into specific transcription regulators in ways that modulate activity of the iron response network that includes Hap43, Sef1, Sfu1 (14, 15, 33). Thus, Mrs4 mutation broadly promotes the induction of iron acquisition pathways even in iron replete conditions as we observed (**Figure 4D**). Thus, *MRS4* mutations may be a key mechanism for a simultaneous increase in activity of multiple regulators. Because inactivating mutations in genes encoding Sfu1, the transcriptional repressor of iron uptake or Yfh1, the iron-sulfur exporter, would only activate a subset of pathways or have other pleiotropic effects on the cell. Unlike LOF or gain-of-function mutations in iron regulators themselves, the *MRS4* mutation did not result in constitutive derepression of the complete iron uptake regulon, as we observed a reduction in expression level of genes involved in the low iron response in iron-replete media and this level of regulation may be beneficial in preventing the accumulation of toxic concentrations of iron and other metals.

In chronic infections, such as those in the CF lung, the host restricts the availability of essential nutrients such as iron via nutritional immunity (34). Host proteins such as lactoferrin, transferrin, and calprotectin sequester iron, making it less accessible to pathogens. Enhanced expression of siderophore acquisition pathways (e.g. SIT1) and surface ferric reductases (e.g. those encoded by *CLUG_02348* and *CLUG_04344*) by *C. lusitaniae* likely provides a significant advantage in acquiring iron from iron sequestering molecules. The increased accumulation of cellular iron may aid cells during fluctuations in iron availability, such as what might occur in the lung environment that experiences cycles of increased inflammation associated with disease exacerbations and inflammation resolution. *Candida* species also have mechanisms to acquire iron from heme which represents approximately 80% of iron within the body. In chronic bacterial infections, evidence suggests that heme utilization is active (35, 36). *Candida* species employ a three-factor system for the acquisition of heme, using the secreted factor Csa2 to bind and ferry heme to the cell surface, where it is brought into the cell by Pga7 and Rbt5. Heme acquisition pathways (*CLUG_4093*, *CLUG_4096*, and *CLUG_4097*) were at significantly higher levels in *MRS4^Q254^*\* than in *MRS4^REF^* (**Figure 5C and D**) in both control and iron chelated conditions. A role for Mrs4 in chronic conditions is an interesting contrast to the importance of Mrs4 function in systemic fungal infections and it is interesting to consider different iron demands and sources in different types of infections (19).

Decreased Mrs4 activity may allow cells to accumulate high levels of iron without jeopardizing mitochondrial function. Iron is highly regulated by all cells due to its reactive properties. *MRS4* mutation may not only enhance iron uptake, but also limit iron concentrations in mitochondria which protects mitochondria from damage. Reduced iron uptake by mitochondria may be key for the accumulation of total cellular iron.

One unique feature of the three CF *C. lusitaniae* infections was that there were no detectable bacterial pathogens as the time of the BAL sample collection. When other *Candida* species are detected in CF, the fungi are usually part of a mixed bacterial-fungal infection. The reason for these intriguing differences is not known and a current area of study. Here, we showed that when compared to *C. albicans* SC5314, *C. lusitaniae* produced significantly less ethanol and more acetate as fermentation products. Ethanol stimulates biofilm formation and virulence factor production in CF bacterial pathogens (37, 38), and thus metabolic differences between fungi may be a differentiating factor. Mutations in *MRS4* may also be an important factor in competing with other microbes for iron. Lastly, *C. albicans* and *C. lusitaniae* differ in the degree to which they stimulate macrophages and thus differences in immune response may play an important role. Future studies will determine how different species persist in chronic infections, the roles specific mutations that are repeatedly under selection, and whether there are common themes across different pathogens. Chronic infection by microbes that are not also human commensals, such as *C. lusitaniae* and *E. dermatiditis*, provide an opportunity to study initial adaptations to the host environment and factors that are most critical for survival and persistence.

## Materials and Methods

### Clinical isolate collection from respiratory samples

Clinical isolates were acquired from sputum and bronchoalveolar lavage (BAL) fluid samples that were plated on YPD (1% yeast extract, 2% peptone, 2% glucose, 1.5% agar) containing gentamycin, blood agar or CHROMagar Candida media then restruck on YPD to obtain single isolates which were then saved in 25% glycerol. *C. lusitaniae* clinical isolates were obtained in accordance with the study protocol approved by Dartmouth Health Institutional Review Board (#22781) using methods described in (5).

### DNA isolation, genome sequence analysis and variant calling

Genomic DNA was extracted from cultures grown in YPD (2% peptone, 1% yeast extract, and 2% glucose) for ∼16 hours; extractions were performed using the MasterPure yeast DNA purification kit (Epicentre). Genomic libraries, for single and pooled isolate DNA, were prepared using the KAPA HyperPrep Kit and sequenced using paired-end 150 bp reads on the Illumina NextSeq500 platform, to a depth of 100-150x coverage per sample as described in Demers *et al.* (5). The pipeline for genome analyses is available in a github repository (https://github.com/stajichlab/PopGenomics_Clusitaniae; doi: 10.5281/zenodo. 7800401). The short read sequences were aligned to a modified version (5) of the *Candida lusitaniae* ATCC 42720 genome (39). The ATCC 42720 genome was altered to remove mitochondrial fragments inserted into the nuclear assembly and the mitochondrial contig (Supercontig_9) was replaced by a complete mitochondrial genome from strain *C. lusitaniae* CBS 6936 (NC_022161.1). The following regions were masked out due to unusually high coverage and likely mitochondrial origin: (Supercontig_1.2:1869020-90 1869184,1664421-1664580; Supercontig_1.3:1076192-1076578,1324802-1324956,1353096-1353260; Supercontig_1.6:126390-126604; Supercontig_1.8:29199-92 29370). Alignments were made using bwa (0.7.17-r1188) (40) and stored as a sorted, aligned read CRAM file with Picard (2.14.1, http://broadinstitute.github.io/picard/) to assign read groups and mark duplicate reads (script 01_align.sh). CRAM files were processed to realign reads using GATK’s RealignerTargetCreator v4.1.8.1 and IndelRealigner following best practices of GATK (41). Each realigned CRAM file was processed with GATK’s HaplotypeCaller (script 02_call_gvcf.sh) followed by joint calling of variants on each Chromosome using GATK’s GenotypeGVCF method script 03_jointGVCF_call_slice.sh). This step also removed low quality variant positions: low quality SNPs were filtered based on mapping quality (score <40), quality by depth (<2 reads), Strand Odds Ratio (SQR>4.0), Fisher Strand Bias (>200), and Read Position Rank Sum Test (<-20). These files were combined to produce a single variant call format (VCF) file of the identified variants to produce list of high quality polymorphisms (script 04_combine_vcf.sh). The quality filtered VCF file containing only variants among the clinical isolates was categorized by SnpEff (5.1) (42) and the ATCC 42720 gene annotation. Genome assemblies of the strains was performed with SPAdes (v3.12.0) (42) after trimming and adaptor cleanup of the reads was performed with AdapatorRemoval (v2.0) (43) and quality trimming with sickle (v1.33) (44). *De novo* assemblies were further screened for vector contamination with vecscreen step of AAFTF v0.3.1 (https://github.com/stajichlab/AAFTF) (doi: <u>10.5281/zenodo.1620526</u>). The metadata for the strains corresponding to the genome sequences for isolates from these three subjects will be described in a Microbial Resource Announcement submitted prior to publication of this work. The raw sequence reads for whole genome sequencing of Subject A, B, and C isolates have been deposited into NCBI sequence read archive under BioProject # PRJNA948351. Details for the isolates and isolate pools from different regions of the lung for Subject A are described in (5). For the analysis of *MRS4* sequences from environmental *C. lusitaniae* strains (**Table S3**) and from clinical isolates obtained from subsequently obtained sputum or stool sample was performed by amplifying *MRS4* using primers DRM031 and DRM032 and sequenced using these primers along with primer ED157 which binds within the *MRS4* sequence (see **Table S4** for primer sequences).

### Strains and mutant construction

Fungal strains and plasmids used in this study are listed in **Table S3**. Fungi were maintained on YPD medium. CRISPR-Cas9 knockout of *MRS4* from clinical isolates was performed using previously described methods (45). For complementation of the reference *MRS4* allele, 5’ UTR and coding region were amplified from ATCC 42740 and assembled into a complementation plasmid with a marker encoding hygromycin resistance (HYG) and 3’ UTR using yeast recombination cloning (46). The complementation plasmid was digested with NotI and KasI resulting in a ∼3500 bp fragment which was transformed into *mrs4Δ* derivatives of representative isolates along with Cas9 and a crRNA targeting the *NAT* marker.

#### Analysis of *Candida auris MRS4* alleles

Clade I (strain B8441), Clade II (B11220), Clade III (B11221), and Clade IV (B11245) were used in the MRS4 sequence comparisons of C. auris isolates. *C. auris MRS4* was amplified from one of two isolates from the CDC Antibiotic Resistance Isolate bank: AR bank #0382 of Clade I (Biosample Accession # SAMN18754596), and AR bank #0383 of Clade III (Biosample Accession # SAMN05379609). *C. auris MRS4* alleles were amplified using primers with 20 base pairs of overlap with *C. lusitaniae MRS4* 5’ UTR and the HYG resistance cassette. *E. dermatitidis MRS4* alleles were synthesized *de novo* by Genscript with the omission of introns, and 20 base pairs of overlap with *C. lusitaniae MRS4* 5’ UTR and the *HYG* resistance cassette. These sequences were then reintroduced into the native locus by complementation cassette by restriction digest, replacing the reference *MRS4* allele with the heterologous sequences. Complementation of heterologous sequences was then performed by the same method as complementation of the reference allele. Primers are listed in **Table S4**.

#### Growth Assays

Unless otherwise stated, strains were grown as 5 ml cultures in YPD overnight (∼16 h), exponential growth aliquots were washed three times and subcultured into experimental medium. For spot titer growth comparisons, cultures were diluted to 1 OD in diH_2_O, diluted 1:10 serially, and spotted in 5 µl volumes on plates. For growth assays in 96 well plates, a starting concentration of 0.005 OD in YPD was used and stated concentrations of BPS, cobalt chloride, or cadmium chloride were added from stock solutions in water. BPS (Sigma CAS# 52746-49-3), cobalt chloride (Sigma CAS# 7791-13-1), and cadmium chloride (Sigma CAS# 654054-66-7) stocks were 100 mM, 100 mM, and 100 µM respectively. Final yield was measured by OD600 at 24 h post-inoculation. For H_2_O_2_ sensitivity, fresh aliquots of 9.8 M H_2_O_2_ were diluted into YPD at the time of inoculation, and growth was measured over the course of 24 h as previously described. Biolog assays were conducted by suspending 0.01 OD of each strain in YNB, and aliquoting 200µl of culture into 192 wells of two proprietary Biolog™ plates PM1 and PM2A with a diverse array of carbon sources. Growth was measured by OD600 over the course of 48 hours, and the final yield is represented in Supplementary Dataset 1.

#### TTC analysis of surface iron reductase activity

After 24 h growth on the indicated medium, a 10 ml solution containing 0.5 mg/ml or 1 mg/ml tetrazolium chloride, 1% molten agar, and 10 mM FeCl_3_ chloride (from 100 mM stock made fresh), if indicated, was carefully pipetted using a 10 ml serological pipette to cover the entirety of the plate. Plates were incubated for the specified time (10 min to 1 h) prior to imaging.

#### Transcriptome analysis of the effects *MRS4* mutation

For RNA isolation, isolates were sub-cultured from overnight cultures into fresh YPD and grown for 6 h which corresponds to cultures in mid-exponential growth phase. Cultures were sub-cultured into six replicate 5 ml cultures of each strain which were incubated at 37C on a rollerdrum. After 5 h of growth, three replicate cultures of each strain were dosed with BPS to a final concentration of 80 µM while the other cultures received water only. Samples were spun down in 15 ml conical tubes, snap-frozen with ethanol and dry ice, and stored for at least 1 h at -80°C. RNA extraction was performed using to MasterPure Yeast RNA Purification kit protocol (Epicentre) according to manufacturer instructions. RNA was submitted to MiGS for RNA Seq analysis. EdgeR was used for normalization and differential gene analysis of raw counts provided by MiGS. RNA-seq data have been submitted to the SRA database: #SUB10993521.

#### qRT-PCR analysis

Culture growth and RNA extraction was performed as described above for the RNA-seq analyses. RNA was DNAse treated with the Turbo DNA-free Kit (Invitrogen). cDNA was synthesized from 500 ng DNAse-treated RNA using the RevertAid H Minus First Strand cDNA Synthesis Kit (Thermo Scientific), following the manufacturer’s instructions for random hexamer primer (IDT) and GC rich template. qRT-PCR was performed on a CFX96 Real-Time System (Bio-Rad), using SsoFast Evergreen Supermix (Bio-Rad) with the primers listed in Table S4. Thermocycler conditions were as follows: 95 °C for 30 s, 40 cycles of 95 °C for 5 s, 65 °C for 3 s and 95 °C for 5 s. Transcripts were normalized to *ACT1* expression.

#### Intracellular iron quantification

Cells from overnight cultures were subcultured into YPD and grown for 5 h before the addition of 80 µM of BPS iron chelator. Samples were taken before adding chelator, and 1 h after iron restriction. Samples were spun down in pre-weighed Eppendorf tubes, washed, and pellets were dried in a vacuum centrifuge for 3 h. Final dry weight was calculated for each pellet, and each was digested with 100µl of 70% HNO_3_. After overnight digestion, samples were heated to 90°C to ensure complete digestion. After dilution with 3.9 ml of diH_2_O, samples were submitted to the Dartmouth Trace Metal Core for ICP-MS analysis of iron content.

#### HPLC Supernatant Analysis

Strains grown in overnight cultures were washed three times in dH_2_O and subcultured in triplicate into YNB minimal media supplemented with 100 mM glucose at a final OD of 0.01. Cultures were allowed to grow for 6 hours at 37° C, then centrifuged at 13,200 RPM for 5 minutes for the separation of insoluble solids and collection of supernatant. Four blank media samples of YNB were also prepared with known concentrations of added carbon sources. One blank was supplemented with 100 mM glucose, 5 mM sodium acetate, 100 mM ethanol, 5 mM sodium citrate, 500 µM sodium lactate, 5 mM sodium succinate, 200 µM sodium pyruvate, and 100 mM glycerol, with two other blanks containing 1:10 and 1:100 dilutions of these carbon sources for the creation of a standard curve within a linear range. The final blank was prepared without any additional carbon sources. For each sample, 400 µl of supernatant was centrifuged at 10,000 RPM for 2.5 minutes through Corning nonsterile nylon 0.22 X-spin filters (#8169), then 20 µl of 10% sulfuric acid as added. Samples were transferred to 2 ml polypropylene snap top microvials for HPLC analysis. Samples were analyzed for levels of various sugars and organic acids utilizing a Shimadzu HPLC (LC-2030) with Biorad Aminex HPX-87H column, LC-20AD pump system, SPD-20AV detector, SIL20AC autosampler, and CTO-20AC column oven.

#### Statistical Analysis

All data were analyzed using Graph Pad Prism 8. The data represent the mean standard deviation of at least three independent experiments with three technical replicates unless stated otherwise. Comparisons were made using a two-tailed, unpaired Student’s T-Test or ANOVA as indicated. One-way ANOVA tests were performed across multiple samples with Tukey’s multiple comparison test for unpaired analyses.

### Code availability

Names of custom codes used for analysis are indicated in where appropriate in above methods. All codes and sequences are available in the indicated github repositories: analysis pipeline and scripts for whole genome genotyping and phylogeny analysis are available at https://github.com/stajichlab/PopGenomics_Clusitaniae. These are archived with Zenodo under DOI: 10.5281/zenodo.7800401. Analysis pipeline for RNA Seq data is available at https://github.com/hoganlab-dartmouth/Clusitaniae_DESeq2.

## Acknowledgements

Research reported in this publication was supported by National Institutes of Health (NIH) grant R01 AI127548 to D.A.H. from the National Institute of Allergy and Infectious Diseases, T32 T32AI007519 (D.R.M). This work was supported by the Cystic Fibrosis Foundation Research Development Program (CFFRDP) STANTO19R0 for the Translational Research Core, P30-DK117469. HL122372 to A.A. from the National Heart, Lung and Blood Institute, and National Institute of General Medical Sciences (NIGMS) of the NIH under award number T32 GM008704 and AI133956 to E.G.D. J.E.S. is a CIFAR Fellow in the program Fungal Kingdom: Threats and Opportunities.

We would like to thank Dr. Daniel Olson (Thayer School of Engineering at Dartmouth) for support for the analysis of supernatant metabolites by HPLC and Kyria Boundry-Mills who provided environmental *C. lusitaniae* strains (Phaff strain collection (UCDFST). Intracellular iron analysis was performed by the Dartmouth Trace Element Core Facility, which was established by grants from the National Institute of Health (NIH) and National Institute of Environmental Health Sciences (NIEHS) Superfund Research Program (P42ES007373). Sequencing services were provided by the Genomics and Molecular Biology Shared Resource Core at Dartmouth (NCI Cancer Center Support Grant 5P30-CA023108). Equipment used was supported by the NIH IDeA award to Dartmouth BioMT P20-GM113132. Analyses were performed using the computational and data storage resources of the University of California-Riverside HPCC funded by grants from the National Science Foundation (NSF) (MRI-1429826) and NIH (1S10OD016290-01A1). Finally, we would like to thank the Dartmouth MCB community which provided routine feedback and provoking questions on this research.

## References

1. Tyrrell J, Callaghan M. 2016. Iron acquisition in the cystic fibrosis lung and potential for novel therapeutic strategies. Microbiology (Reading) 162:191–205.

2. Gifford AH, Moulton LA, Dorman DB, Olbina G, Westerman M, Parker HW, Stanton BA, O’Toole GA. 2012. Iron homeostasis during cystic fibrosis pulmonary exacerbation. Clin Transl Sci 5:368–73.

3. Vermilyea DM, Crocker AW, Gifford AH, Hogan DA. 2021. Calprotectin-Mediated Zinc Chelation Inhibits Pseudomonas aeruginosa Protease Activity in Cystic Fibrosis Sputum. J Bacteriol 203:e0010021.

4. Kim SH, Clark ST, Surendra A, Copeland JK, Wang PW, Ammar R, Collins C, Tullis DE, Nislow C, Hwang DM, Guttman DS, Cowen LE. 2015. Global Analysis of the Fungal Microbiome in Cystic Fibrosis Patients Reveals Loss of Function of the Transcriptional Repressor Nrg1 as a Mechanism of Pathogen Adaptation. PLoS Pathog 11:e1005308.

5. Demers EG, Biermann AR, Masonjones S, Crocker AW, Ashare A, Stajich JE, Hogan DA. 2018. Evolution of drug resistance in an antifungal-naive chronic *Candida lusitaniae* infection. Proc Natl Acad Sci U S A 115:12040–12045.

6. Ross BS, Lofgren LA, Ashare A, Stajich JE, Cramer RA. 2021. *Aspergillus fumigatus* In-Host HOG Pathway Mutation for Cystic Fibrosis Lung Microenvironment Persistence. mBio 12:e0215321.

7. Hogan DA, Willger SD, Dolben EL, Hampton TH, Stanton BA, Morrison HG, Sogin ML, Czum J, Ashare A. 2016. Analysis of Lung Microbiota in Bronchoalveolar Lavage, Protected Brush and Sputum Samples from Subjects with Mild-To-Moderate Cystic Fibrosis Lung Disease. PLoS One 11:e0149998.

8. Young LY, Hull CM, Heitman J. 2003. Disruption of ergosterol biosynthesis confers resistance to amphotericin B in Candida lusitaniae. Antimicrob Agents Chemother 47:2717–24.

9. Desnos-Ollivier M, Moquet O, Chouaki T, Guerin AM, Dromer F. 2011. Development of echinocandin resistance in Clavispora lusitaniae during caspofungin treatment. J Clin Microbiol 49:2304–6.

10. Asner SA, Giulieri S, Diezi M, Marchetti O, Sanglard D. 2015. Acquired Multidrug Antifungal Resistance in Candida lusitaniae during Therapy. Antimicrob Agents Chemother 59:7715–22.

11. Du H, Bing J, Hu T, Ennis CL, Nobile CJ, Huang G. 2020. *Candida auris*: Epidemiology, biology, antifungal resistance, and virulence. PLoS Pathog 16:e1008921.

12. WHO. 2022. WHO fungal priority pathogens list to guide research, development and public health action., Geneva.

13. Zhang Y, Lyver ER, Knight SA, Pain D, Lesuisse E, Dancis A. 2006. Mrs3p, Mrs4p, and frataxin provide iron for Fe-S cluster synthesis in mitochondria. J Biol Chem 281:22493–502.

14. Muhlenhoff U, Stadler JA, Richhardt N, Seubert A, Eickhorst T, Schweyen RJ, Lill R, Wiesenberger G. 2003. A specific role of the yeast mitochondrial carriers MRS3/4p in mitochondrial iron acquisition under iron-limiting conditions. J Biol Chem 278:40612–20.

15. Li L, Kaplan J. 2004. A mitochondrial-vacuolar signaling pathway in yeast that affects iron and copper metabolism. J Biol Chem 279:33653–61.

16. Froschauer EM, Schweyen RJ, Wiesenberger G. 2009. The yeast mitochondrial carrier proteins Mrs3p/Mrs4p mediate iron transport across the inner mitochondrial membrane. Biochim Biophys Acta 1788:1044–50.

17. Foury F, Roganti T. 2002. Deletion of the mitochondrial carrier genes *MRS3* and *MRS4* suppresses mitochondrial iron accumulation in a yeast frataxin-deficient strain. J Biol Chem 277:24475–83.

18. Gupta M, Outten CE. 2020. Iron-sulfur cluster signaling: The common thread in fungal iron regulation. Curr Opin Chem Biol 55:189–201.

19. Xu N, Dong Y, Cheng X, Yu Q, Qian K, Mao J, Jia C, Ding X, Zhang B, Chen Y, Zhang B, Xing L, Li M. 2014. Cellular iron homeostasis mediated by the Mrs4-Ccc1-Smf3 pathway is essential for mitochondrial function, morphogenesis and virulence in *Candida albicans*. Biochim Biophys Acta 1843:629–39.

20. Xu N, Cheng X, Yu Q, Zhang B, Ding X, Xing L, Li M. 2012. Identification and functional characterization of mitochondrial carrier Mrs4 in *Candida albicans*. FEMS Yeast Res 12:844–58.

21. Yates CM, Filippis I, Kelley LA, Sternberg MJ. 2014. SuSPect: enhanced prediction of single amino acid variant (SAV) phenotype using network features. J Mol Biol 426:2692–701.

22. Choi Y, Do E, Hu G, Caza M, Horianopoulos LC, Kronstad JW, Jung WH. 2020. Involvement of Mrs3/4 in Mitochondrial Iron Transport and Metabolism in *Cryptococcus neoformans*. J Microbiol Biotechnol 30:1142–1148.

23. Reedy JL, Floyd AM, Heitman J. 2009. Mechanistic plasticity of sexual reproduction and meiosis in the *Candida* pathogenic species complex. Curr Biol 19:891–9.

24. Nyhus KJ, Ozaki LS, Jacobson ES. 2002. Role of mitochondrial carrier protein Mrs3/4 in iron acquisition and oxidative stress resistance of *Cryptococcus neoformans*. Med Mycol 40:581–91.

25. Chen C, Pande K, French SD, Tuch BB, Noble SM. 2011. An iron homeostasis regulatory circuit with reciprocal roles in *Candida albicans* commensalism and pathogenesis. Cell Host Microbe 10:118–35.

26. Skrahina V, Brock M, Hube B, Brunke S. 2017. *Candida albicans* Hap43 Domains Are Required under Iron Starvation but Not Excess. Front Microbiol 8:2388.

27. Weissman Z, Shemer R, Kornitzer D. 2002. Deletion of the copper transporter CaCCC2 reveals two distinct pathways for iron acquisition in *Candida albicans*. Mol Microbiol 44:1551–60.

28. Alvarez-Ortega C, Harwood CS. 2007. Responses of *Pseudomonas aeruginosa* to low oxygen indicate that growth in the cystic fibrosis lung is by aerobic respiration. Mol Microbiol 65:153–165.

29. Lockhart SR, Etienne KA, Vallabhaneni S, Farooqi J, Chowdhary A, Govender NP, Colombo AL, Calvo B, Cuomo CA, Desjardins CA, Berkow EL, Castanheira M, Magobo RE, Jabeen K, Asghar RJ, Meis JF, Jackson B, Chiller T, Litvintseva AP. 2017. Simultaneous Emergence of Multidrug-Resistant *Candida auris* on 3 Continents Confirmed by Whole-Genome Sequencing and Epidemiological Analyses. Clin Infect Dis 64:134–140.

30. Munoz JF, Gade L, Chow NA, Loparev VN, Juieng P, Berkow EL, Farrer RA, Litvintseva AP, Cuomo CA. 2018. Genomic insights into multidrug-resistance, mating and virulence in Candida auris and related emerging species. Nat Commun 9:5346.

31. Kurbessoian T, Murante D, Crocker A, Hogan DA, Stajich J. 2022. In host evolution of *Exophiala dermatitidis* in cystic fibrosis lung micro-environment. bioRxiv 2022.09.23.509114.

32. Murad AM, Leng P, Straffon M, Wishart J, Macaskill S, MacCallum D, Schnell N, Talibi D, Marechal D, Tekaia F, d’Enfert C, Gaillardin C, Odds FC, Brown AJ. 2001. NRG1 represses yeast-hypha morphogenesis and hypha-specific gene expression in Candida albicans. Embo J 20:4742–52.

33. Wofford JD, Lindahl PA. 2015. Mitochondrial Iron-Sulfur Cluster Activity and Cytosolic Iron Regulate Iron Traffic in *Saccharomyces cerevisiae*. J Biol Chem 290:26968–26977.

34. Fourie R, Kuloyo OO, Mochochoko BM, Albertyn J, Pohl CH. 2018. Iron at the Centre of *Candida albicans* Interactions. Front Cell Infect Microbiol 8:185.

35. Nguyen AT, O’Neill MJ, Watts AM, Robson CL, Lamont IL, Wilks A, Oglesby-Sherrouse AG. 2014. Adaptation of iron homeostasis pathways by a *Pseudomonas aeruginosa* pyoverdine mutant in the cystic fibrosis lung. J Bacteriol 196:2265–76.

36. Skaar EP, Humayun M, Bae T, DeBord KL, Schneewind O. 2004. Iron-source preference of *Staphylococcus aureus* infections. Science 305:1626–8.

37. Chen AI, Dolben EF, Okegbe C, Harty CE, Golub Y, Thao S, Ha DG, Willger SD, O’Toole GA, Harwood CS, Dietrich LE, Hogan DA. 2014. Candida albicans ethanol stimulates Pseudomonas aeruginosa WspR-controlled biofilm formation as part of a cyclic relationship involving phenazines. PLoS Pathog 10:e1004480.

38. Doing G, Koeppen K, Occipinti P, Harty CE, Hogan DA. 2020. Conditional antagonism in co-cultures of Pseudomonas aeruginosa and Candida albicans: An intersection of ethanol and phosphate signaling distilled from dual-seq transcriptomics. PLoS Genet 16:e1008783.

39. Butler G, Rasmussen MD, Lin MF, Santos MA, Sakthikumar S, Munro CA, Rheinbay E, Grabherr M, Forche A, Reedy JL, Agrafioti I, Arnaud MB, Bates S, Brown AJ, Brunke S, Costanzo MC, Fitzpatrick DA, de Groot PW, Harris D, Hoyer LL, Hube B, Klis FM, Kodira C, Lennard N, Logue ME, Martin R, Neiman AM, Nikolaou E, Quail MA, Quinn J, Santos MC, Schmitzberger FF, Sherlock G, Shah P, Silverstein KA, Skrzypek MS, Soll D, Staggs R, Stansfield I, Stumpf MP, Sudbery PE, Srikantha T, Zeng Q, Berman J, Berriman M, Heitman J, Gow NA, Lorenz MC, Birren BW, Kellis M, et al. 2009. Evolution of pathogenicity and sexual reproduction in eight *Candida* genomes. Nature 459:657–62.

40. Li H, Durbin R. 2009. Fast and accurate short read alignment with Burrows-Wheeler transform. Bioinformatics 25:1754–60.

41. Van der Auwera GA, Carneiro MO, Hartl C, Poplin R, Del Angel G, Levy-Moonshine A, Jordan T, Shakir K, Roazen D, Thibault J, Banks E, Garimella KV, Altshuler D, Gabriel S, DePristo MA. 2013. From FastQ data to high confidence variant calls: the Genome Analysis Toolkit best practices pipeline. Curr Protoc Bioinformatics 43:11 10 1-11 10 33.

42. Cingolani P, Platts A, Wang le L, Coon M, Nguyen T, Wang L, Land SJ, Lu X, Ruden DM. 2012. A program for annotating and predicting the effects of single nucleotide polymorphisms, SnpEff: SNPs in the genome of *Drosophila melanogaster* strain w1118; iso-2; iso-3. Fly (Austin) 6:80–92.

43. Schubert M, Lindgreen S, Orlando L. 2016. AdapterRemoval v2: rapid adapter trimming, identification, and read merging. BMC Res Notes 9:88.

44. N. Joshi JF. 2011. Sickle: A sliding-window, adaptive, quality-based trimming tool for FastQ files (Version 1.33), Available at https://github.com/najoshi/sickle.

45. Grahl N, Demers EG, Crocker AW, Hogan DA. 2017. Use of RNA-Protein Complexes for Genome Editing in Non-albicans *Candida* Species. mSphere 2.

46. Shanks RM, Caiazza NC, Hinsa SM, Toutain CM, O’Toole GA. 2006. Saccharomyces cerevisiae-based molecular tool kit for manipulation of genes from gram-negative bacteria. Appl Environ Microbiol 72:5027–36.

