## Supplementary material for "Mrs4 loss of function in fungi during adaptation to the cystic fibrosis lung": Figures S1-S7 and Tables S1-S4

```
CLUSTAL O(1.2.4) multiple sequence alignment

UT8656          ----MTLMNSPD---AVPGEEFDYEALPPNYGLGHNMLAGAFAGIAEHTVMYPVDLMKTR     53
S288C           MVENSSSNNSTRPIPAIPMDLPDYEALPTHAPLYHQLIAGAFAGIMEHSVMFPIDALKTR     60
SC5314          ------MEHQLQFLPKDS-VEIDYEALPDDASLAAHLTAGALAGIMEHTVMFPIDSIKTR     53
ATCC42740       ------MADHPHQLGADT-LEVDYEALPEDASIIAHLSAGAFAGIMEHTVMYPIDSIKTR     53
B11220          ------MADHPHMLS-EP-IEIDYEALPEDASIAAHLSAGAFAGIMEHTVMFPIDSIKTR     52
                        .             ****** .  :  :: ***:*** **:**:*:* :***

UT8656          MQIINPSAGGLYTGLSHAVSTIYRLEGLRTLWRGVTSVIVGAGPAHAVYFGTYEMVKELA     113
S288C           IQSANAKS-LSAKNMLSQISHISTSEGTLALWKGVQSVILGAGPAHAVYFGTYEFCKKNL     119
SC5314          MQMNLSNS-EISRGLLKSLSKISSTEGFYALWKGVSSVVLGAGPAHAIYFSVFESTKTFL     112
ATCC42740       MQIIASGQ-ASSRSVISSISRISSSEGAYALWRGVSSVVMGAGPAHAVYFSVFEATKTML     112
B11220          MQMIASGQ-PLSKSVIGSISKISSSEGAYALWRGVSSVVLGAGPAHAVYFSVFEATKTML     111
                :*           .:   :* *   **  :**:** **::*******:**..:*  *

UT8656          GTNST---------DGKHHPFAAAASGAAATITSDALMNPFDVIKQRMQVHGST------     158
S288C           IDSSD---------TQTHHPFKTAISGACATTASDALMNPFDTIKQRIQLNTS-------     163
SC5314          VNRLTNSPHSNRIVTDENHPLIASCAGITGTTASDALMTPFDMLKQRMQANAAYQDGK--     170
ATCC42740       VNRLTNSS-STKIVTDESHPLIASGAGIAATIASDALMTPFDVLKQRMQVGNEAKA----     167
B11220          VNRLTNSR-STKIVTDESHPLIASGAGIAATVASDALMTPFDVLKQRMQLGTSTADSKLT     170
                                  **: :: :*  .* :*****.*** :***:*

UT8656          ---YRSLTHCAREIFRTEGFSAFYVSYPTTLCMTVPFTATQFMAYESLSTIMNPKKEYDP     215
S288C           ----ASVWQTTKQIYQSEGLAAFYYSYPTTLVMNIPFAAFNFVIYESSTKFLNPSNEYNP     219
SC5314          -STSVRLFKLASDIYKAEGLSAFYISYPTTLLTNIPFAALNFGFYEYSSSLLNPSHVYNP     229
ATCC42740       --SSVKLLSTALSIYKTEGASAFFISYPTTLFTNIPFAALNFGFYEYCSSLLNPENTYNP     225
B11220          QSKSARLVKTAAQIYSKEGLSAFFISYPTTLFTNIPFAALNFGFYEYSSKLLNPENKYNP     230
                      :   : .*:  ** :**: ******  .:**:* :*  **  :.::**.: *:*

UT8656          ITHCVAGGLAGAFAAGITTPLDVIKTLLQTRGLSQ--KDEIRNVRGLFHAASIIKREFGW     273
S288C           LIHCLCGSISGSTCAAITTPLDCIKTVLQIRGSQTVSLEIMRKADTFSKAASAIYQVYGW     279
SC5314          YLHCVSGGIAGGIAAALTTPFDCIKTVLQTKGISQ--NQNFRHVTGFKSAAVALLKQEGA     287
ATCC42740       YYHCVSGGIAGGIAAALTNPFDCIKTALQTRGIST--NESLRNVNGFSSAARALYRHGGF     283
B11220          YYHCISGGIAGGIAAALTNPLDCIKTALQTRGVSQ--NESLRTINGFVSAAKALYKQGGA     288
                  **:.*.::*. .*.:*.*:* *** ** :* .    : :*    :  **  : :  *

UT8656          SGFMRGWRPRIISTMPSTAICWSSYEMAKAYFKRTLREEHEAQMSKL  320
S288C           KGFWRGWKPRIVANMPATAISWTAYECAKHFLMTY------------  314
SC5314          KAFWKGLKPRVIFNIPSTAISWTAYEMCKEVLIRGKGL---------  325
ATCC42740       GAFMRGLKPRIIFNVPSTAISWTAYEMAKEVLLRD------------  318
B11220          GAFTRGLKPRIIFNVPSTAISWTAYEMAKEVLLRD------------  323
                 .* :* :**:: .:*:***.*::** .*  :
```

**Figure S1. Non-synonymous SNPs occur in highly conserved regions of Mrs4 orthologs.** Clustal Omega(Sievers et al., 2011)generated alignment of representative *MRS4* peptide sequences from *E. dermatitidis (UT6856), S. cerevisiae (S288C), C. albicans* (SC5314), *C. lusitaniae (ATCC 42740),* and *C. auris (B11220).* Missense and nonsense mutations with functional effects are highlighted: C. lusitaniae G138 (light purple), A147 (dark purple), A235 (red), and Q254 (cyan). The residue that corresponds to *E. dermatitidis* E40 (brown) is also shown.

Synonymous SNPs

| | DH2383 | ATCC | DH3115 | DH3116 | DH3117 | DH3118 | DH3119 | DH3120 | DH3121 | DH3122 |
|---|---|---|---|---|---|---|---|---|---|---|
| DH2383 | 0 | | | | | | | | | |
| ATCC | 0 | 0 | | | | | | | | |
| DH3115 | 1 | 1 | 0 | | | | | | | |
| DH3116 | 1 | 1 | 0 | 0 | | | | | | |
| DH3117 | 0 | 0 | 1 | 1 | 0 | | | | | |
| DH3118 | 1 | 1 | 0 | 0 | 1 | 0 | | | | |
| DH3119 | 5 | 5 | 6 | 6 | 5 | 6 | 0 | | | |
| DH3120 | 5 | 5 | 6 | 6 | 5 | 6 | 0 | 0 | | |
| DH3121 | 6 | 6 | 6 | 7 | 6 | 6 | 4 | 4 | 0 | |
| DH3122 | 6 | 6 | 7 | 7 | 7 | 7 | 5 | 5 | 5 | 0 |

**Figure S2. The *MRS4* sequences of environmental, reference, and acute infection isolates of *C. lusitaniae* differ by only a small number of synonymous SNPs.** Comparison of synonymous SNPs in the *MRS4* sequences encoding the “reference” amino acid sequence. *MRS4* alleles of *C. lusitaniae* clinical strains (DH2383 and ATCC 42740) and environmental isolates (see Table S3) differ only by the number of synonymous mutations indicated in the matrix.

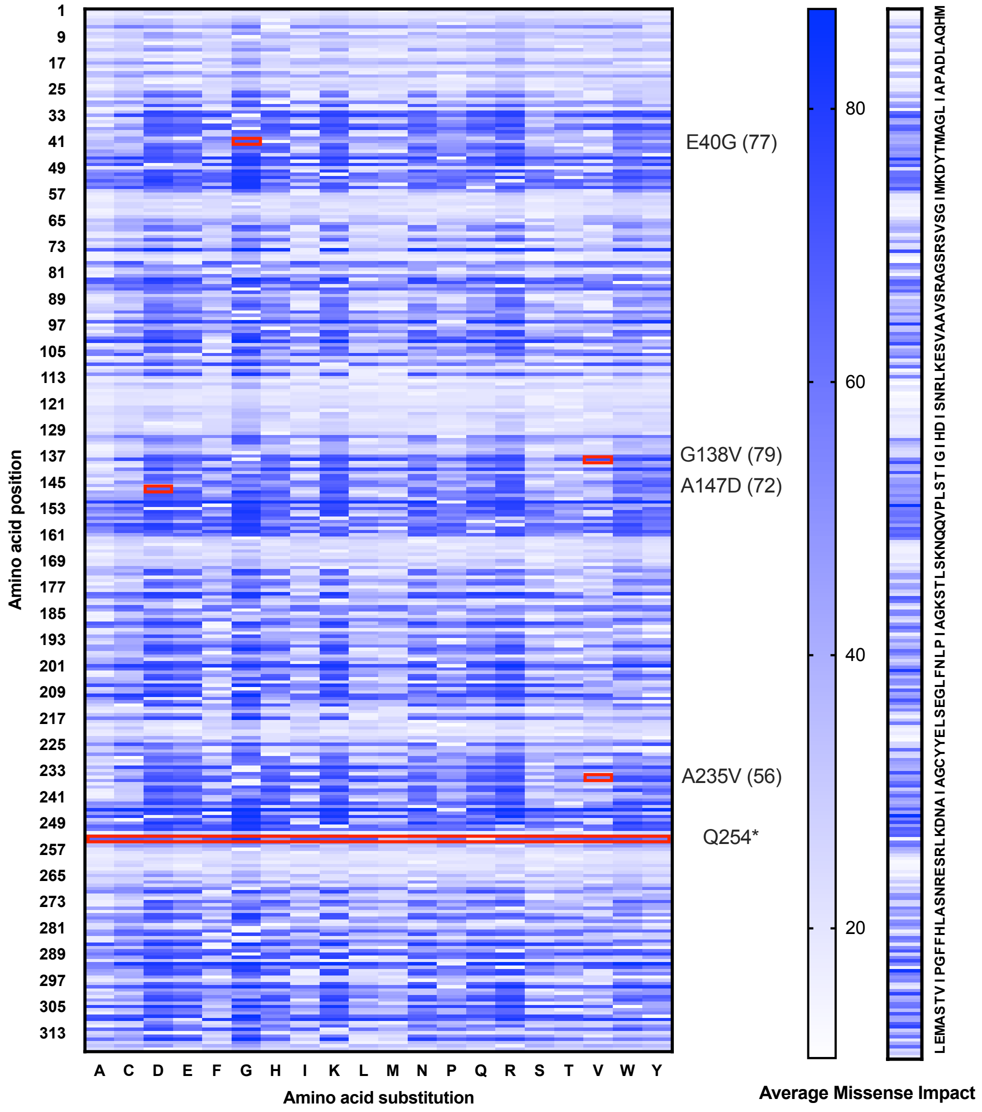


Yates CM, Filippis I, Kelley LA & Sternberg MJE (2014) SuSPect: Enhanced prediction of single amino acid variant (SAV) phenotype using network features. *Journal of Molecular Biology*. In press. http://dx.doi.org/10.1016/j.jmb.2014.04.026

**Figure S3. Non-synonymous SNPs occur in predicted high-severity impact regions**. SuSPect (Yates, Filippis, Kelley, & Sternberg, 2014) mutational analysis heatmaps indicate that the predicted severity of each possible particular amino acid substitution at a given position and the average impact of all possible amino acid substitutions on a scale from 0 (least severe) to 100 (most severe). Higher areas of impact coincide with predicted transmembrane alpha helix regions (see **Fig. 1A**). *C. lusitaniae MRS4* mutations are highlighted in red with their predicted impact score in parentheses (except for the Q254* mutation). The predicted effects of the *E. dermatiditis MRS4* substitution, which leads to an E40G substitution, is also shown.

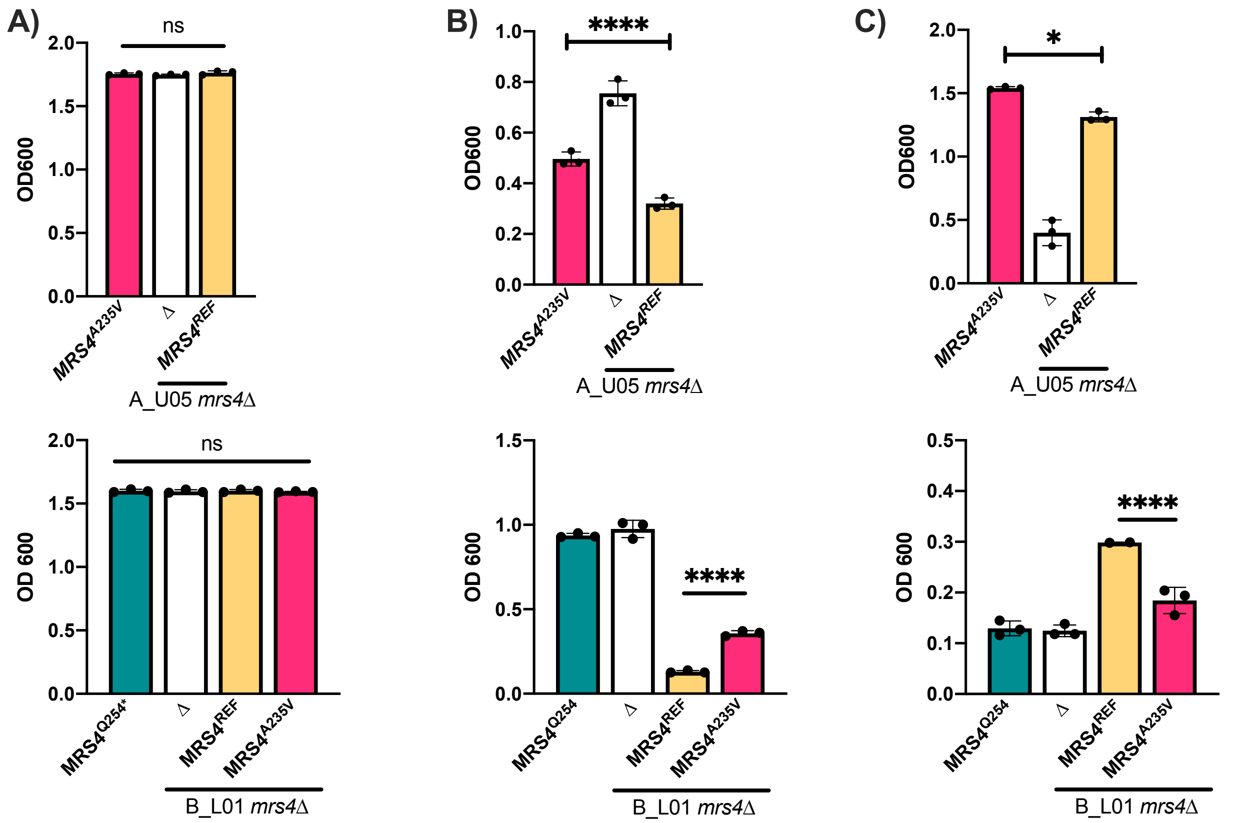


**Figure S4. The A_U10D strain background masks the Mrs4-dependent phenotype of cadmium sensitivity. A)** Effects of *MRS4*$^{A235V}$ on growth in YPD in the A_U05 (top) and B_L01 (bottom) backgrounds. The *mrs4*Δ, and *mrs4*Δ+*MRS4*$^{REF}$ are shown for both strains and *mrs4*Δ+MRS4A235V in the B10D background. Growth for the same strains as in panel A in 2.5 mM $CoCl_3$ **B)** and 12.5 mM $CdCl_2$ **C)** are shown. Data shown in panel C (top) is also shown as part of Fig. 2F.

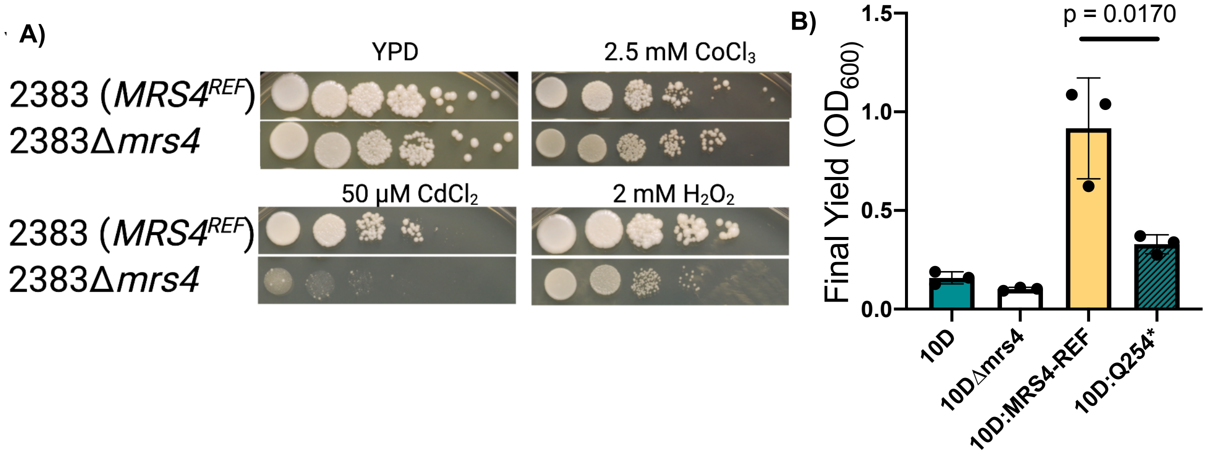


**Figure S5. Deletion of *MRS4* from strain 2383 with the *MRS4*$^{REF}$ allele has the predicted Mrs4 loss-of-function phenotypes. A)** An *mrs4*Δ strain, constructed in DH2383, which has *MRS4*$^{REF}$ sequence, and the DH2383 parent strain spotted in a dilution series on YPD plates supplemented with cobalt, cadmium, and hydrogen peroxide as described in the Materials and Methods for 48 h before imaging. **B)** B_L01 parent, *mrs4*Δ*,* and complemented strains with the *MRS4*$^{REF}$ or native *MRS*$^{Q254*}$ allele were grown in a 96-well plate format in YPD containing 1 mM $H_2O_2$ for 24 h at 37˚. Final yield was measured as absorbance at 600 nm on a Synergy Neo2 plate reader.

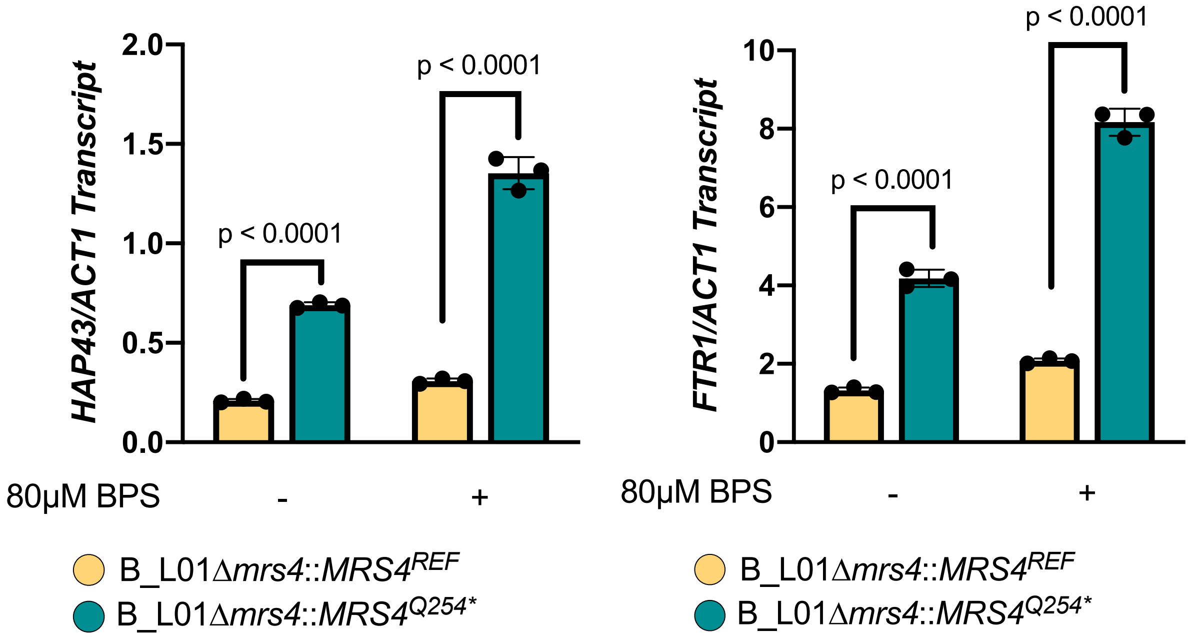


**Figure S6. Quantitative RT-PCR analysis of *HAP43* and *FTR1* in the absence and presence of Mrs4 activity in iron replete and iron-chelated conditions.** Transcripts encoding the iron-regulating transcription factor Hap43 and high-affinity iron transporter Ftr1 were measured in B_L01 *mrs4*Δ strains complemented with $MRS4^{REF}$ or native $MRS^{Q254*}$ in cells grown in YPD for 6 h, or in YPD for 5 h followed by 1 h growth after the addition of 80μM BPS.

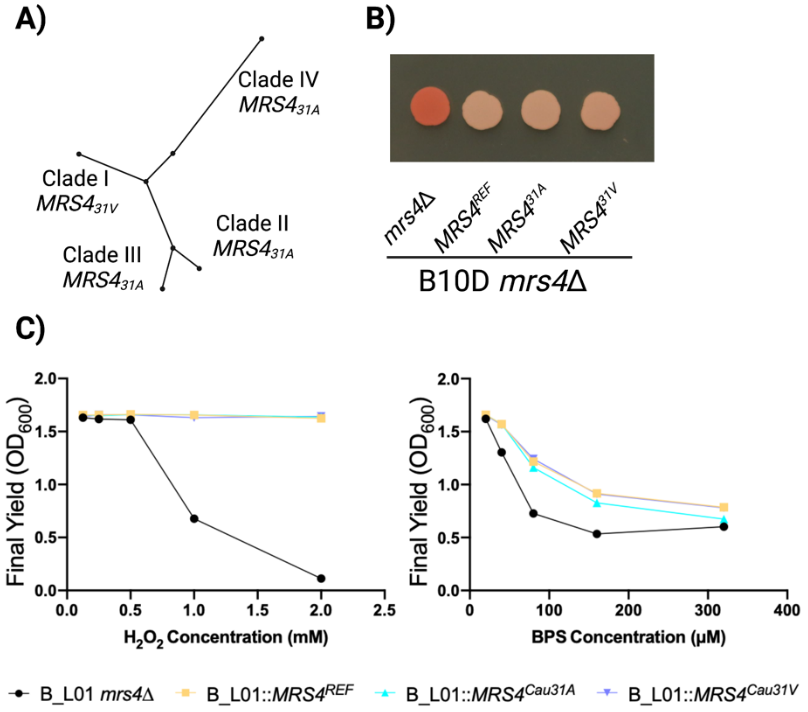


**Figure S7. The *C. auris* clade I ortholog of *MRS4* differs from other clades by a single amino acid change that does not affect Mrs4 function. A)** *C. auris* clade distribution of *MRS4* alleles, with phylogenetic tree based on (Du et al., 2020) **B)** TTC reduction phenotypes of *C. auris MRS4* alleles heterologously expressed in the B_L01 background, spotted on YPD plates and grown for 24 h before TTC overlay. **C)** Growth of *C. auris* alleles in BPS and $H_2O_2$.

**Table S1:** Loci with heterogeneity in encoded amino acid sequences within more than one CF-associated *C. lusitaniae* population.

| ***Variable loci in Subj. A and B (4)*** | ***Variable loci in Subj. A and C (7)*** | ***Variable loci in Subj. B and C (8)*** |
|---|---|---|
| CLUG_03891 | CLUG_03989 | CLUG_03981 |
| CLUG_00379 | CLUG_01165 | CLUG_03274 |
| CLUG_03468 | CLUG_05802 | CLUG_01977 |
| CLUG_02919 | CLUG_02244 | CLUG_02910 |
| | CLUG_00052 | CLUG_04108 |
| | CLUG_05353 | CLUG_01201 |
| | CLUG_03194 | CLUG_01123 |
| | | CLUG_05462 |

**Table S2: Mrs4 loss of function does not alter secondary carbon metabolite secretion in *C. lusitaniae.*** B_L01 derivatives B_L01::*MRS4*$^{REF}$ and B_L01::*MRS4*$^{Q254*}$, and *C. albicans* SC5314 with single and double knockouts of *MRS4,* were cultured overnight and sub-cultured into YNB minimal media with 100mM glucose as a primary carbon source. Supernatants were collected and analyzed by HPLC for consumption of glucose, and production of major secondary metabolites. Molar carbon ratios were calculated for consumed glucose and citrate to produced acetate, ethanol, and glycerol, relative to the base media.

| | *C. lusitaniae* B_L01 | | *C. albicans* SC5314 | |
|---|---|---|---|---|
| Substrates (mM) | *MRS4*$^{REF}$ | *MRS4Q254** | *MRS4/ MRS4* | *mrs4* Δ/Δ |
| Glucose utilized | 43.9 | 27.5 | 63 | 59.6 |
| Citrate utilized | 2.6 | 2.37 | 2.50 | 1.90 |
| **Average Carbon Consumed** | **279** | **179** | **392** | **369** |
| Acetate produced | 41.4 | 23.9 | 3.38 | 2.56 |
| Ethanol produced | 13.4 | 5.14 | 24.9 | 29.3 |
| Glycerol produced | nd | nd | 0.08 | 0.02 |
| **Average Carbon Produced** | **110** | **58.1** | **56.9** | **63.8** |
| **Carbon produced: Carbon utilized** | **0.39** | **0.32** | **0.15** | **0.17** |

**Table S3. Strains and plasmids**

| Strain | Strain background | Marker | Lab # | Source |
|---|---|---|---|---|
| ***C. lusitaniae*** | | | | |
| ATCC 42740 | | | DH2387 | (Butler et al., 2009) |
| DH2383(RSY284/ CL6936) | | | DH2383 | (Reedy, Floyd, & Heitn |
| *mrs4Δ* | DH2383 | NatR | DH4093 | This study |
| A_U05 | | | DH3087 | Dartmouth Health |
| B_L01 | | | DH3703 | Dartmouth Health |
| *mrs4* | " | NatR | DH4080 | This study |
| *mrs4::MRS4*$^{REF}$ | " | HygB-R | DH4081 | This study |
| *mrs4::MRS4*$^{Q254*}$ | " | HygB-R | DH4082 | This study |
| *mrs4::MRS4*$^{Cau-31V}$ *(C. auris)* | " | HygB-R | DH4083 | This study |
| *mrs4::MRS4*$^{Cau-31A}$ *(C. auris)* | " | HygB-R | DH4084 | This study |
| *mrs4::MRS4*$^{Ed-40G}$ *(E. dermatiditis)* | " | HygB-R | DH4085 | This study |
| *Mrs4::MRS4*$^{Ed-40E}$ *(E. dermatiditis)* | " | HygB-R | DH4086 | This study |
| A_U05 *mrs4Δ* | A_U05 | NatR | DH4087 | This study |
| *mrs4Δ::MRS4*$^{REF}$ | A_U05 | HygB-R | DH4088 | This study |
| B_L04 | | | DH3706 | Dartmouth Health |
| *mrs4Δ* | B_L04 | NatR | DH4089 | This study |
| *mrs4Δ::MRS4*$^{REF}$ | B_L04 | HygB-R | DH4090 | This study |
| C_M06 | | | DH3708 | Dartmouth Health |
| *mrs4Δ* | C_M06 | NatR | DH4091 | This study |
| *mrs4Δ::MRS4*$^{REF}$ | C_M06 | HygB-R | DH4092 | This study |
| A_S22 | Sp2 isolate from Subj. A | | | |
| A_S23 | " | | | |
| A_S24 | " | | | |
| A_S25 | " | | | |
| B_S17 | Sp4 from subject B | | | |
| B_S18 | " | | | |
| 61-4 | Environ. | | DH3115 | PYCC** |
| 71-129 | Environ. | | DH3116 | PYCC** |
| 76-31 | Environ. | | DH3117 | PYCC** |
| 79-1 | Environ. | | DH3118 | PYCC** |
| 80-11 | Environ. | | DH3119 | PYCC** |
| 80-12 | Environ. | | DH3120 | PYCC** |

| | | | | |
|---|---|---|---|---|
| 82-606.2 | Environ. | | DH3121 | PYCC** |
| 16-4994.55 | Environ. | | DH3122 | PYCC** |
| ATCC 42720 | Clinical | | DH2387 | (Butler et al., 2009) |
| Acute 1A | Acute | | DH3769 | This study |
| AR0398 | Unknown | | DH2785 | CDC panel |
| 1H | Clinical, ascending colon, *MRS4*$^{A147D}$ | | DH4021 | This study |
| 6B | Clinical, transverse colon, *MRS4*$^{A147D}$ | | DH4023 | This study |
| 6D | Clinical, transverse colon, *MRS4*$^{A147D}$ | | DH4025 | This study |
| 11B | Clinical, descending colon, *MRS4*$^{A147D}$ | | DH4031 | This study |
| 11G | Clinical, descending colon, *MRS4*$^{A147D}$ | | DH4036 | This study |
| 11H | Clinical, descending colon, *MRS4*$^{A147D}$ | | DH4037 | This study |
| ***Candida albicans*** | | | | |
| SC5314 | | | DH35 | |
| *mrs4* | SC5314 | NatR | DH4094 | This study |
| *mrs4*Δ/Δ | SC5314 | NatR | DH4095 | This study |
| **Plasmids** | | | | |
| pDRM03 | *pMQ30-MRS4REF-HygR* | | DH4096 | |
| pDRM06 | *pMQ30-MRS4Q254*-HygR* | | DH4097 | |
| pDRM10 | *pMQ30-MRS4-*$^{31A}$*(C. auris)* | | DH4098 | |
| pDRM13 | *pMQ30-MRS3-*$^{31V}$(*C. auris*) | | DH4099 | |
| pDRM14 | *pMQ30-MRS440G* (*E. dermatiditis*)*-HygR* | | DH4100 | |
| pDRM15 | *pMQ30-MRS440E* (*E. dermatiditis*)*-HygR* | | DH4101 | |

**Table S4. Primers**

| Primer Label | Sequence | Descriptor |
|---|---|---|
| ED149 | CTTTGCCTCACATGGGCACTC | CLUG_02526 LF FWD |
| ED150 | gtattctgggcctccatgtcGGTTGGATGGTAGAATCAGAAGAAG | CLUG_02526 LF REV |
| ED151 | gaatgctggtcgctatactgGATTGCGGACATGTGTATTGGTTG | CLUG_02526 RF FWD |
| ED152 | CACTAGATCATGGAGTATGCGG | CLUG_02526 RF REV |
| ED153 | CTTCTTCTGATTCTACCATCCAACCgacatggaggcccagaatac | CLUG_02526 + NAT FWD |
| ED154 | CAACCAATACACATGTCCGCAATCcagtatagcgaccagcattc | CLUG_02526 + NAT REV |
| ED155 | GATATGGGTTGTGGACAATCTCCG | CLUG_02526 NEST FWD |
| ED156 | CTCTTCTAACTCATGCCTGATCTCG | CLUG_02526 NEST REV |
| ED157 | CATCTGCTATCTGTTGCTG | CLUG_02526 Sanger Sequencing |
| ED179 | GATTAAGTTGGGTAACGCCAGGCGCCCTTTGCCTCACATGGGCACTC | CLUG_02526 LF Fwd + pMQ30 overlap |
| ED180 | CAAGCACTATACGACGTCAGGTCTAGAGTAGACAATGTGCCTATACTTG | CLUG_02526 Rev + Hyg overlap |
| ED181 | CAAGTATAGGCACATTGTCTACTCTAGACCTGACGTCGTATAGTGCTTG | Hyg FWD + CLUG_02526 overlap |
| ED182 | GTTTCGTATGCAATGTAATGGATAGAAAGGGCCTCGTGATACGC | Hyg Rev + CLUG_02526 overlap |
| ED183 | GCGTATCACGAGGCCCTTTCTATCCATTACATTGCATACGAAAC | CLUG_02526 RF Fwd + Hyg overlap |
| ED184 | GTATGTTGTGTGGAATTGTGAGCGGCCGCCTCTTCTAACTCATGCCTGATC | CLUG_02526 RF rev + pMQ30 overlap |
| DRM019 | CTTGTATGAGGCGGTCTTTC | pDRM03 Sequencing |
| DRM020 | CAACTTGGGGCTGACACAC | pDRM03 Sequencing |
| DRM021 | GACAGAGGGTGCCAGTGC | pDRM03 Sequencing |
| DRM022 | CATGTAATAAACGGATACGGCA | pDRM03 Sequencing |
| DRM023 | GATGTATTTACCAACGCCATC | pDRM03 Sequencing |
| DRM024 | CTTTCGAAACATGGATAACGAG | pDRM03 Sequencing |
| DRM031 | ggcattaaagaagggcaagg | *mrs4* External 5' check |
| DRM032 | gactcatctcctggcatgaa | *mrs4* External 3' check |
| DRM041 | GATTAAGTTGGGTAACGCCAATGGCCGACCATCCCCATATG | C. auris MRS4 Fwd + CLUG_02526 LF overlap |
| DRM042 | AAGCACTATACGACGTCAGGCTAGTCTCTCAACAACACCTCCTTGG | C. auris MRS4 Rev + Hyg overlap |
| DRM045 | TCTGATTCTACCATCCAACCATGACTCTAATGAACAGCCCGGATG | E. derm MRS4 Fwd + CLUG_02526 LF overlap |
| DRM047 | AAGCACTATACGACGTCAGGTCAGAGCTTGCTCATTTGAGCC | E. derm MRS4 Rev + Hyg overlap |
| DRM073 | CTTCTGATTCTACCATCCAACC | pDRM014/015 sequencing |
| DRM074 | CCGGTCGCTCACACATTG | pDRM014/015 sequencing |
| DRM075 | CAGAGCTTGCTCATTTGAGCC | pDRM014/015 sequencing |

| | | |
|---|---|---|
| DRM076 | CACGGCGATTTGCTGGTC | pDRM014/015 sequencing |
| DRM106 | CCGCGTCAGATGCTTTGATGCAAATTAAAAATAGTTTACGCAAGTC | SNR52-CaMRS4proto linker Rev |
| DRM107 | CATCAAAGCATCTGACGCGGGTTTTAGAGCTAGAAATAGCAAGTTAAA | Calbicans-MRS4 Protolinker-sgRNA scaffold Fwd |
| DRM108 | AGAAGCATAAAGAAACAAGCAATTCGTGGCTCAGACAAGGACACAGATAATACTTAAATTAATAATACAATACTATCATATTTTCCCAGTCACGACGTT | Calbicans-MRS4 linked-NAT KO Cassette Fwd |
| DRM109 | TAGTAATGAAACAAAAACGTTATCTGAAATTATTTACAAATTAGAAAAATAAATACCAAGTAGTATGATGTCTATGAAAAGTGGAATTGTGAGCGGATA | Calbicans-MRS4 linked-NAT KO Cassette Rev |
| DRM110 | GAAGTGTGCGTGTGTGAGC | C. albicans MRS4 KO 5' Check |
| DRM111 | GCGCCCAATACCACAGAAG | C. albicans MRS4 KO 3' Check |

Butler, G., Rasmussen, M. D., Lin, M. F., Santos, M. A., Sakthikumar, S., Munro, C. A., . . . Cuomo, C. A. (2009). Evolution of pathogenicity and sexual reproduction in eight *Candida* genomes. *Nature, 459*(7247), 657-662. doi:10.1038/nature08064

Du, H., Bing, J., Hu, T., Ennis, C. L., Nobile, C. J., & Huang, G. (2020). Candida auris: Epidemiology, biology, antifungal resistance, and virulence. *PLoS Pathog, 16*(10), e1008921. doi:10.1371/journal.ppat.1008921

Reedy, J. L., Floyd, A. M., & Heitman, J. (2009). Mechanistic plasticity of sexual reproduction and meiosis in the Candida pathogenic species complex. *Curr Biol, 19*(11), 891-899. doi:10.1016/j.cub.2009.04.058

Sievers, F., Wilm, A., Dineen, D., Gibson, T. J., Karplus, K., Li, W., . . . Higgins, D. G. (2011). Fast, scalable generation of high-quality protein multiple sequence alignments using Clustal Omega. *Mol Syst Biol, 7*, 539. doi:10.1038/msb.2011.75

Yates, C. M., Filippis, I., Kelley, L. A., & Sternberg, M. J. (2014). SuSPect: enhanced prediction of single amino acid variant (SAV) phenotype using network features. *J Mol Biol, 426*(14), 2692-2701. doi:10.1016/j.jmb.2014.04.026
